# Drag-induced directionality switching of kinesin-5 Cin8 revealed by cluster-motility analysis

**DOI:** 10.1101/2020.01.24.918714

**Authors:** Himanshu Pandey, Emanuel Reithmann, Alina Goldstein-Levitin, Jawdat Al-Bassam, Erwin Frey, Larisa Gheber

## Abstract

Directed active motion of motor proteins is a vital process in virtually all eukaryotic cells. Nearly a decade ago, the discovery of directionality switching of mitotic kinesin-5 motors challenged the long-standing paradigm that individual kinesin motors are characterized by an intrinsic directionality. While several kinesin motors have now been shown to exhibit context-dependent directionality that can be altered under diverse experimental conditions, the underlying mechanism remains unknown. Here, we studied clustering-induced directionality switching of the mitotic kinesin-5 Cin8, using a fluorescence-based single-molecule motility assay combined with biophysical theory. Based on the detailed characterization of the motility of single motors and clusters of Cin8, we developed a predictive molecular model, that quantitatively agrees with experimental data. This combined approach allowed us to quantify the response of Cin8 motors to external forces as well as the interactions between Cin8 motors, and thereby develop a detailed understanding of the molecular mechanism underlying directionality switching. The main insight is that directionality switching is caused by a single feature of Cin8: an asymmetric response of active motion to forces that oppose motion, here referred to as drag. This general mechanism explains why bidirectional motor proteins are capable of reversing direction in response to seemingly unrelated experimental factors including clustering, changes in the ionic strength of the buffer, increased motor density and molecular crowding, and in motility assays.

**Significance Statement:** Kinesin-5 motor proteins perform essential functions in chromosome segregation during mitotic cell division. Surprisingly, several kinesin-5 motors have the ability to reverse directionality under different experimental conditions, which contradicts the long-standing paradigm that individual kinesin motors are characterized by an intrinsic directionality. The mechanism underlying this ability to switch directionality has remained elusive. Here, we combine fluorescence-based motility assays and theoretical modeling to analyze cluster-size-dependent motility of the bidirectional kinesin-5 Cin8. Our results show that bidirectional motors can switch directionality because they exhibit an asymmetric response of active motion to drag. This mechanism explains multiple seemingly unrelated experimental factors that have been shown to cause directionality switching of kinesin motors.

## Introduction

The bipolar and tetrameric kinesin-5 motors perform essential functions in mitotic spindle dynamics by crosslinking and sliding antiparallel spindle microtubules (MTs) apart (1-3). In kinesin-5 motors the catalytic domain is located at the amino-terminus. Since they share this attribute with all known plus-end-directed kinesins, they were previously believed to be exclusively plus-end directed. Remarkably, recent studies showed that several kinesin-5 motor proteins, which are in fact minus-end directed at the single-molecule level, can switch directionality under various experimental conditions. Thus far, three bidirectional kinesin-5 motors have been identified (4-7): Cin8 and Kip1 in *Saccharomyces cerevisiae* and Cut7 in *Schizosaccharomyces pombe*. Their *in-vitro* behavior is puzzling: Single Cin8 and Kip1 motors are minus-end directed in high ionic strength but switch directionality under low-ionic-strength conditions (5,7,8), as well as in multi-motor gliding assays (5-7). Cin8 was also shown to switch directionality when engaged in sliding antiparallel MTs apart (6,7), as a function of the density of surface-bound motors that interact with MTs in multi-motor gliding assays (9), and when it forms clusters on single MTs (10). Cut7 is minus-end directed in single-molecule experiments and multi-motor MT gliding assays (4,11), but switches directionality as a function of crowding of either motor or non-motor proteins on MTs (12). Notably, two kinesin-14 motors were recently demonstrated to be bidirectional (13,14), indicating that switchable directionality is more common in the kinesin superfamily than was previously appreciated. In fact, the plethora of observations related to directionality switching may indicate that bidirectionality is of physiological importance, a notion that is also supported by a recent theoretical study (15).

Presently, however, there is no mechanistic explanation for all these intriguing observations and the remarkable ability of the kinesin-5 motors to switch directionality remains unexplained. Here we study directionality switching due to clustering of motor proteins (10), employing a combined experimental and theoretical approach. We developed a single-molecule fluorescence-based method of analysis that enabled us to characterize the relationship between Cin8 cluster size and directionality. We found that while single Cin8 motors show minus-end-directed motion, clusters of Cin8 containing two and more Cin8 motors are more likely to display plus-end-directed motion. The motion of single Cin8 motors exhibits a large diffusive component, which is markedly reduced for Cin8 pairs and higher-order clusters. By relating predictions derived from a computational model to experimental data for the differential cluster-size-dependent motion of Cin8, we gain mechanistic insights into Cin8 motility and interactions. More specifically, we find that the motion of individual motors exhibits an active, directed component in both directions along the MT, together with a diffusive component akin to Brownian motion. Importantly, our analysis shows that the plus- and minus-end-directed active components of Cin8 motion respond *asymmetrically* to forces that *oppose* this motion (the latter are referred to here as “*drag*”), and that motors in clusters interact via weak attractive forces. The interplay of these various factors leads to clustering-induced directionality switching of Cin8 from minus-to plus-end-directed motility. Our study yields detailed insights into the mechanism underlying the directionality switching of the bidirectional kinesin motor Cin8. The very same mechanism explains not only directionality switching due to motor clustering, but also provides an understanding of directionality switching under diverse, previously reported conditions, such as changes in the ionic strength or the motor density in the motility assay, and molecular crowding with other motor and non-motor proteins. Thus, we provide a general mechanism for directionality switching, which advances our understanding of the origin of bidirectional motion of motor proteins.

## Results

### Motile clusters containing two Cin8 molecules can switch directionality

We have previously reported that clustering of Cin8 motors on MTs is one of the conditions that can switch its directionality from fast minus-end- to slow plus-end-directed movement (10). However, this analysis did not provide quantitative information on the relationship between directionality and the number of motors in a cluster, which is essential for unraveling the underlying mechanism. Hence, to establish a basis for the analysis of the link between Cin8 clustering and the preferred direction of motility, we first developed an experimental strategy to measure the *cluster size*, i.e. the number of Cin8 molecules in a cluster. By following the fluorescence intensity of Cin8-GFP as a function of time, we observed single photobleaching events of ∼50 arbitrary units (a.u.), most likely stemming from bleaching of a single GFP molecule (Fig. 1A). Since single Cin8 motors are tetramers containing four identical subunits, the maximal fluorescence intensity of a single Cin8 motor containing four GFP molecules is likely to be ⁓200 a.u. Based on this method of quantification, the population of Cin8 motors or clusters was divided into three categories according to the number of Cin8 molecules in each: (i) individual or “*single”* tetrameric Cin8 molecules (intensities < 200 a.u.), (ii) “*pairs”* of Cin8 molecules i.e. dimers of *single* tetrameric Cin8 molecules (intensities 200-400 a.u.), and (iii) higher “*oligomers”* of Cin8 (intensities > 400 a.u.) (Fig. SA1 in the Supporting Information). Pairs and higher oligomers of Cin8 are also referred to as “clusters” henceforth. “Cin8 motors” refers to moving Cin8 particles of unspecified cluster-size. Further critical assessment of the experimental approach used to determine the Cin8 cluster size is given in section A2.1 of the Supporting Information.

**Figure 1:**
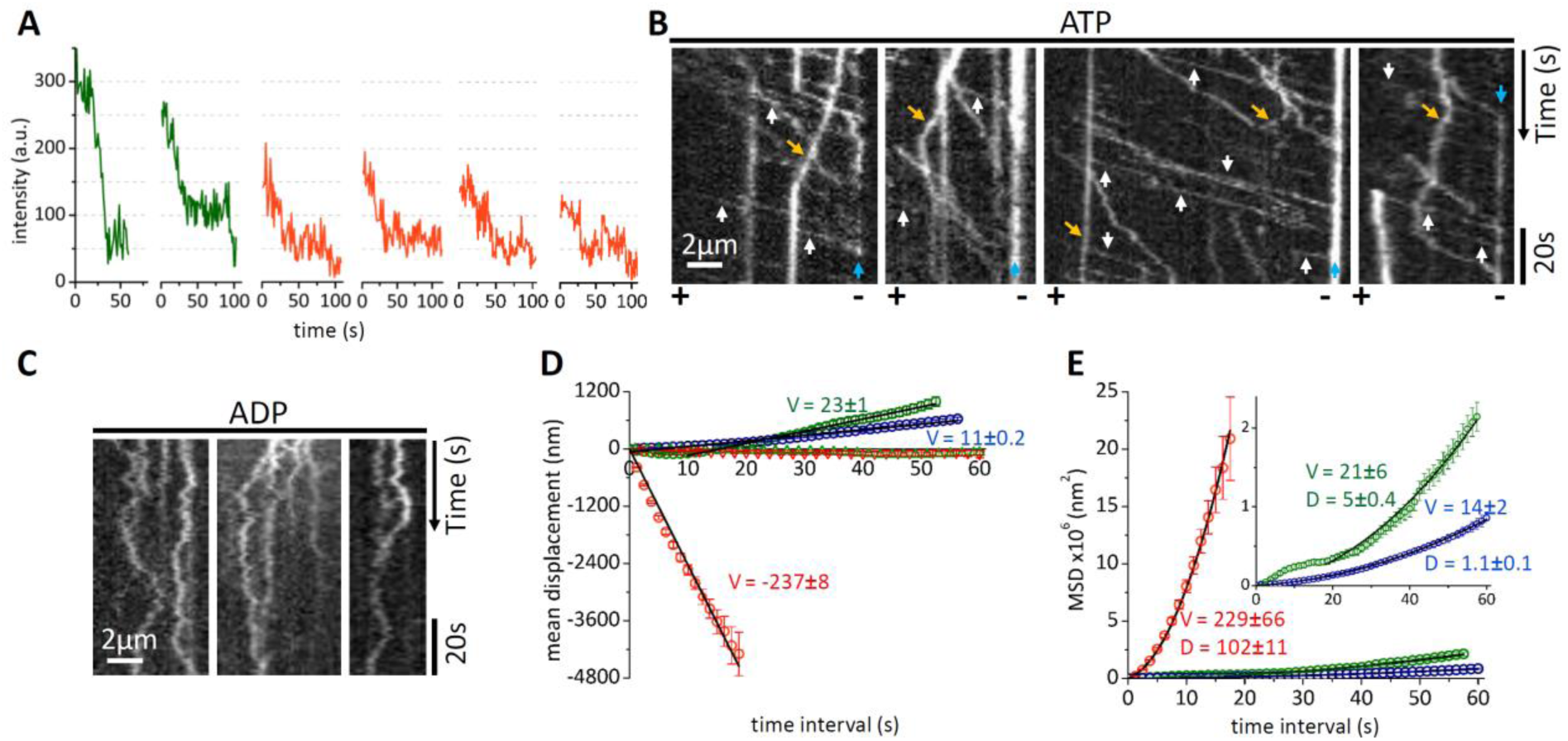
Motility of Cin8 motors of different size. **(A)** Photobleaching profile of fluorescent Cin8-GFP motors. Single photobleaching steps, each likely representing the photobleaching of one GFP, lead to a drop in fluorescence intensity of ⁓50 a.u. (arbitrary units). The red and green traces represent single motors and pairs of Cin8, respectively. **(B)** Representative kymographs of Cin8-GFP motility along single MTs in the presence of 1 mM ATP. White arrows: fast-moving Cin8 motors; orange arrows: plus-end-directed Cin8 motors; blue arrows: clustering of Cin8 at the minus end of the MTs (10). Polarity of the MTs is indicated at the bottom. **(C)** Representative kymographs of Cin8-GFP motility along single MTs in the presence of 1 mM ADP**. (D, E)** Mean displacements (MD) **(D)** and mean-square displacements (MSD) **(E)** of single Cin8 molecules (red), pairs of Cin8 (green) and higher oligomers (blue) in the presence of 1 mM ATP (circles) or ADP (triangles). The black solid lines are linear (MD = vt) (MD) or statistically weighted, second-order polynomial fits (MSD = v^2^t^2^+2Dt) (MSD). Values of the mean velocity (V, nm/s) and the diffusion coefficient (D, nm^2^/s (x10^3^) are indicated. Error bars represent SEM. Negative values of velocity represent motility in the minus-end direction. In **(E)**, the inset shows a zoomed-in plot depicting the MSD curves for Cin8 pairs (green) and oligomers (blue).

To study the relation between Cin8 cluster size and motility, we performed experiments at saturating ATP concentration and high ionic strength (Fig. 1B), where Cin8 motors were previously shown to be mostly minus-end directed and fast-moving relative to kinesin-5 motors in higher eukaryotes (6,7,10,16). In accordance with previous reports (6,7,10,16), we observed that ∼80% of the total motile Cin8 population was fast-moving and minus-end directed, and that almost all these trajectories fell into the category of single tetrameric Cin8 motors (Fig. 1A, SA2a (left panel) and Table 1). The remaining 20% comprise clusters of Cin8, which exhibit increased incidences of plus-end-directed and bidirectional trajectories (Fig. 1B, SA2a (middle and right panels) and Table 1). Importantly, we found that MT-bound clusters of Cin8 move along MTs (Fig. SA2a) and undergo splitting events and mergers (Fig. SA3), indicating that these are active motors rather than non-specific Cin8 aggregates. Moreover, clustering occurred regardless of the nucleotide state, indicating that it is an intrinsic property of Cin8. Thus, motile clusters of Cin8 were observed both under high and low ATP concentrations and in the presence of ADP (Fig. 1B, C; SA2; SA4 and SA5). Experiments carried out in the presence of ATP clearly showed that the cluster size correlated with the propensity for plus-end-directed motion. While only 3% of single-molecule trajectories were on average plus-end directed, 40% and 70% of the trajectories of pairs and oligomers of Cin8, respectively, exhibited net movement towards the plus end (Fig. SA2a). Notably, in each cluster-size category, the velocity of minus-end-directed motion was considerably faster than that of plus-end-directed motility (Fig. SA2a).

**Table 1:**
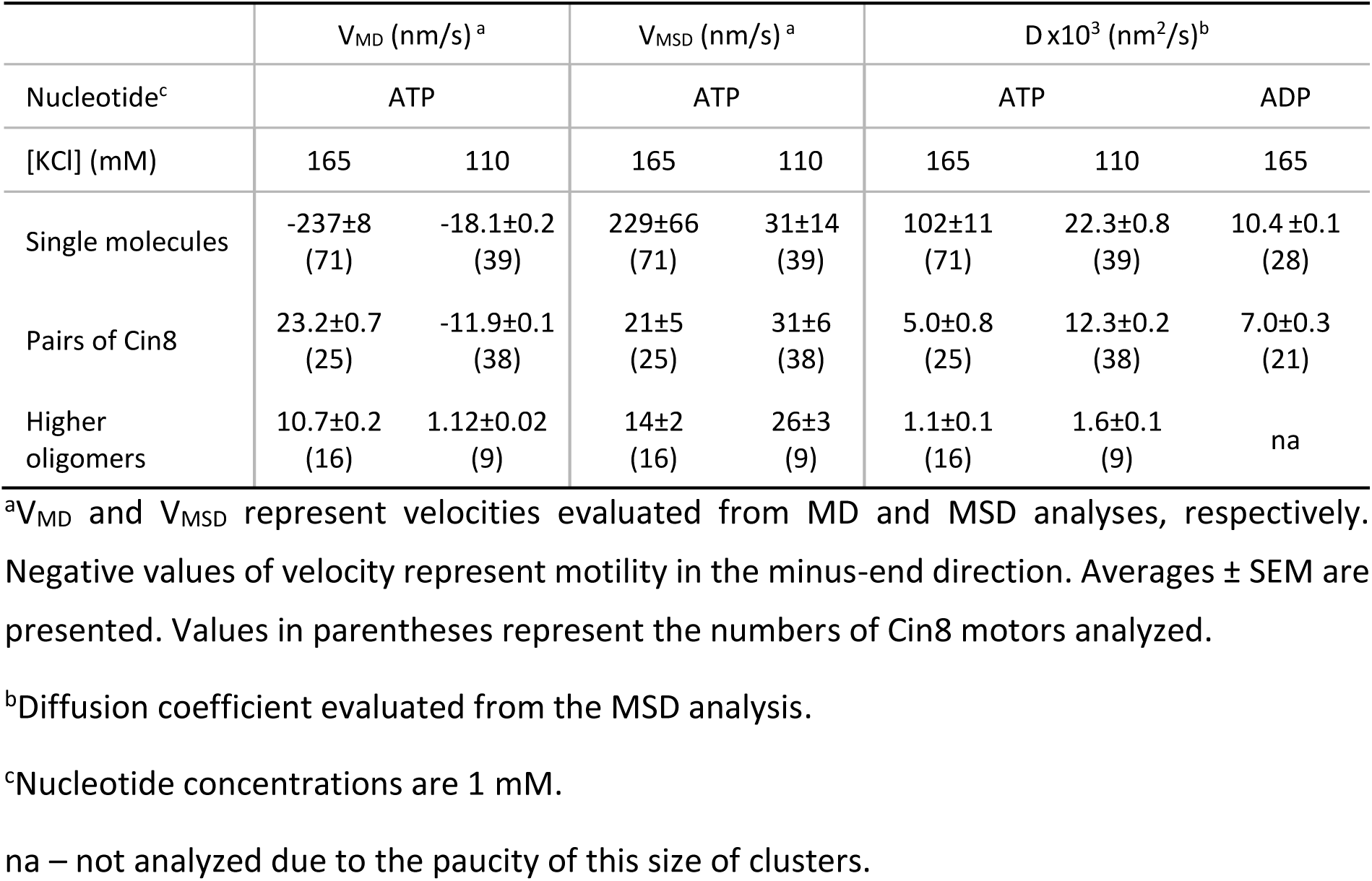
MD and MSD analysis of the motility of Cin8-GFP motors

In summary, as a basis for further analyses, we developed an experimental approach that enabled us to assess the number of Cin8 molecules in a moving cluster. This in turn allowed us to quantitatively show that the preference for slow, plus-end-directed motion correlates with increasing size of Cin8 clusters, and that pairs of Cin8 motors can switch directionality.

### Individual Cin8 motors exhibit a large diffusive component

To better understand the implications of Cin8 clustering, we next sought to characterize the motion of single Cin8 molecules and Cin8 clusters. As central quantities for motility, we used the average displacement velocity and the diffusion coefficient, obtained from an analysis of the mean displacement (MD) and mean-squared displacement (MSD) (Fig. 1D, E, SA6; Table 1). This quantitative analysis further supported the observation that single Cin8 motors are fast and minus-end directed (v = −237 nm/s), while pairs and clusters of Cin8, on average, are slow and plus-end directed with velocities of v = 23 nm/s and v = 11 nm/s, respectively (Fig. 1D and Table 1). In the presence of ATP, the motor dynamics deviates from diffusive behavior, as the slope of the log-log plot of the MSD against time is greater than one. This result confirms that single Cin8 molecules and Cin8 clusters exhibit active motion (Fig. SA6a, b) (8,17). In contrast, and as expected, we observed only diffusive motion in the presence of ADP (Fig. SA6c, d) (8,17).

Next, we focused on the motility of single Cin8 molecules in order to establish a basis for a mechanistic model of Cin8 motion. In addition to the large MSD velocity reported above, single Cin8 molecules also exhibited an exceptionally high effective diffusion coefficient of D ≈ 102 · 10^3^ nm^2^/s in the presence of ATP (Fig. 1E and Table 1). This value may relate to a large bidirectional component in the motion of Cin8, which could arise from active bidirectional stepping, Brownian motion or a combination of both. In the presence of ADP, the diffusion coefficient was substantially lower with a value of D ≈ 10 · 10^3^ nm^2^/s (Table 1). The considerable decrease in the diffusion coefficient in the presence of ADP relative to ATP suggests at first sight that single Cin8 motors may indeed exhibit active bidirectional stepping that leads to additional effective diffusive behavior, which vanishes in the absence of ATP. There is, however, an additional counteracting effect. The average dwell times of ADP-bound Cin8 (Table SA1) are longer, indicating an increased affinity of Cin8 for MTs under this condition, which, in turn, is expected to reduce the diffusion coefficient. To discriminate between the different possible origins of the large diffusion coefficient, we estimated the potential ATP turnover rate required to obtain the observed diffusion coefficient by active bidirectional stepping alone, and obtained a value of 3000 ATPs/sec (see section B3 of the Supporting Information for a detailed computation). Such a high turnover rate exceeds known values for other kinesins by at least one order of magnitude (18-20). Moreover, the diffusion coefficient measured for Cin8 in the presence of ATP shows a value that is typical for molecules that exhibit an ATP-independent diffusive component of motion along MTs (21-23). Thus, it is likely that the largest contribution to the diffusion coefficient of individual Cin8 motors originates from Brownian motion. Here, it is important to note that this argument does not exclude the presence of an additional, but significantly smaller, contribution of ATP-dependent (active) bidirectional motion to the diffusion coefficient.

Overall, we conclude from the statistical analysis of the ensemble of stochastic motor trajectories that single Cin8 molecules show an active, ATP-dependent component, leading to minus-end-directed motion of single motors and plus-end-directed motion of clusters. Single Cin8 motors exhibit a strong diffusive component, which is largely attributable to Brownian motion.

### Cin8 motors are subject to weak attractive interactions that reduce cluster diffusion

Based on this understanding of the motion of single Cin8 motors, we asked whether groups of Cin8 display markedly different behavior. By performing an MSD analysis as detailed in the previous section, we found that clusters of Cin8 motors indeed exhibited a substantially decreased diffusion coefficient as compared to single Cin8 motors (Fig. 1E and Table 1). For example, the diffusion coefficient for pairs of Cin8 was approximately twenty times smaller than that of single motors (Table 1). Based on our observations, we hypothesized that weak attractive interactions between Cin8 tetramers cause clustering of motors, and might also be the underlying cause of the reduced diffusion coefficients of clusters.

To test these hypotheses, we formulated a computational model designed specifically to predict the consequences of such weak interactions for the motion and clustering of Cin8 motors and, ultimately, for their directionality. This model comprises two main modules, one for the diffusive and one for the active components of the motions of the motors. To derive such a model from experimental data and test its predictions, we proceeded in two steps. First, we quantified the model parameters related to Cin8 interactions by establishing a functional relation between cluster diffusion and attractive interactions (while neglecting active motion). This approach is justified as, based on the above considerations, the diffusive component of motion mainly reflects the impact of Brownian motion. In the next step, elaborated in the following section, we added a module that accounts for active motion and its response to motor interactions, thereby establishing a mechanistic link between clustering and direction switching.

In the module that captures the diffusive behavior, we considered each single, non-interacting Cin8 molecules as a random walker moving bidirectionally on the MT lattice with equal hopping rates to the plus and minus ends; the hopping rate was obtained from the measured diffusion coefficient for single Cin8 molecules (Fig. 1E and Table 1). We assumed that Cin8 molecules interact via a weak and constant force *F*_int_ if they are within a given interaction range *R* of each other (Fig. 2A). As dictated by equilibrium statistical physics (detailed balance), this force must exert an effect on the hopping rates of interacting Cin8 motors through a Boltzmann factor *e*^−Δ*E*/*k*^*_B_^T^*, where Δ*E* = *F*_int_ *a* is the energy required to move two motors a distance *a* = 8.4 nm (the length of a tubulin dimer) apart (see section B1 of the Supporting Information for a technical description of the stochastic model and the numerical implementation of the stochastic particle dynamics using a Gillespie algorithm (24)). An *in silico* parameter scan over different values for the magnitude of the attractive force and the interaction range yielded two crucial insights. First, our simulations strongly suggested an interaction range of approximately *R* = 25 nm, since smaller or larger values led to unstable clusters or very large clusters, respectively, which is at variance with experimental observations (see Fig. SB5 and section B2 of the Supporting Information for the corresponding simulation results with different values of *R*). Notably, this interaction range is in accordance with the length of kinesin-5 motors (approximately 80 nm (25-28)). Second, we observed that increasing the attractive force *F*_int_ between Cin8 molecules indeed causes a strong decrease in the diffusion coefficient of Cin8 pairs, as shown in Fig. 2B. This monotonic decrease originates from the fact that the interaction between two motors effectively acts as a load when the motors move away from each other. As a consequence, increasing Cin8 interaction strength decreases the diffusive motion of a Cin8 pair or cluster. From the relationship between the attractive force *F*_int_ and the diffusion coefficient *D* we inferred that the measured diffusion coefficient of *D* = 5 · 10^3^ nm^2^/s for a pair of Cin8 corresponds to an attractive force of *F*_int_ = 1.4 pN, (Fig. 2B, dashed line).^1^ This value is close to the stall force for Cin8 (≈ 1.5 pN) observed in motility assay experiments (9).

**Figure 2:**
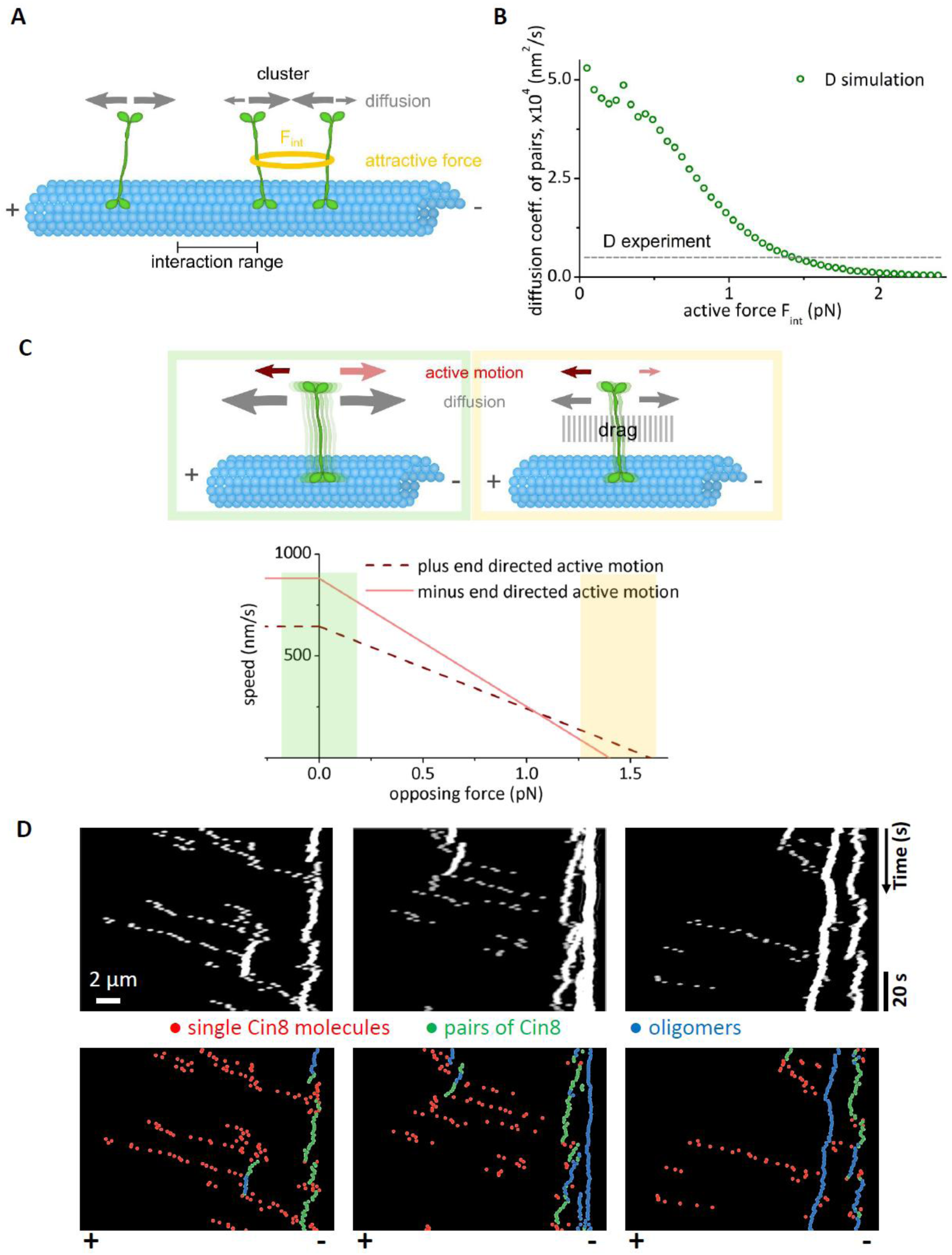
Computational model for collective Cin8 motion. **(A)** The diffusive module of the computational model simulates Cin8 motors as random walkers with symmetric dynamics (gray arrows) due to Brownian motion. Motors attract each other if they are within the interaction range *R*. The attractive force *Fint* (illustrated as a yellow ring) is assumed to be constant for intermotor distances less than *R* and zero otherwise. Attractive interactions affect motor dynamics (detailed balance) as described in the main text and section B1 of the Supporting Information. The model was implemented by means of a one-dimensional lattice gas as detailed in section B1 of the Supporting information. **(B)** The diffusion coefficient of pairs of Cin8 in simulations (circles) decreased with increasing attractive forces *F* between motors. Attractive forces of 1.4 pN between motors result in a diffusion coefficient for Cin8 pairs that corresponds to that identified *in vitro* (dashed line; see also Table 1). **(C, Top)** Overall Cin8 dynamics in the computational model is composed of bidirectional active motion (red/pink arrows in the upper panels) and a diffusive component (gray arrows; illustrated by motor blurring in the upper panels) as described in **(A)**. In the absence of an opposing force, the minus-end-directed active component is larger than the plus-end-directed component (left panel, red arrows). Both the active and diffusive components of the model are affected by forces that oppose this motion, which are here referred to as ‘drag’ (gray shaded area in the upper right panel). Drag suppresses Brownian motion exponentially, and minus-end-directed active motion (pink) is more strongly impeded by drag than is plus-end-directed active motion (red), such that the overall directionality switches under large drag. **(C, Bottom)** The assumed force-velocity relation used in simulations, with green shading corresponding to low drag (as illustrated in the upper left panel) and yellow shading corresponding to high drag (as illustrated in the upper right panel). The arrows in **(A)** and **(C)** are for illustrative purposes only; their sizes are not proportional to the model parameters. **(D)** Simulated kymographs exhibit directionality switching due to motor clustering. Cin8 clusters were biased towards the plus end, while single motors were biased towards the minus end. In the upper row, simulation data was convoluted with a point-spread function to obtain images comparable to the experimental data, where bright traces represent clusters (see section B9 of the Supporting information for details of the convolution method). The bottom row shows the same kymographs, but here single Cin8 motors (red), pairs of Cin8 (green) and larger clusters with more than two Cin8 molecules (blue) are depicted in different colors. Polarity of the MTs is indicated at the bottom. Identical parameters were used in the three simulations in **(D)**. Please refer to section B2 and Table SB1 the Supporting Information for a detailed description and a complete list of the parameters employed in **(B)** and **(D)**.

We next asked whether, with these inferred parameters, the theoretical model can predict the experimentally observed clustering statistics. Indeed, we found that upon varying the Cin8 concentration, the changes in the cluster-size distribution predicted by simulations closely matched those observed experimentally (Fig. 2C, D and SB7).

### A drag asymmetry of active motion explains directionality switching of Cin8 clusters

After successfully establishing the role of attractive interactions in the diffusive motion of Cin8 and cluster formation, we next investigated whether attractive interactions might be related to clustering-induced directionality switching. To address this question, we extended the model to account for active motion, based on the following assumptions (Fig. 2C). First, motivated by the difference in diffusion coefficients of single Cin8 motors in the presence of ATP and ADP observed *in vitro* (Table 1), we hypothesized that single Cin8 motors show bidirectional active motion towards both ends (red/pink arrows in Fig. 2C). As single Cin8 molecules were found to be biased towards the minus-end in the experiments, such active bidirectional motion must also be biased in this direction in the absence of external forces. Second, as a basic mechanism that could potentially switch the directionality of clusters (Fig. 1D and SA2a), we hypothesized that motors that are moving towards the plus and minus ends, respectively, respond differently to forces that oppose this motion (collectively termed “drag”). Specifically, we assumed that drag suppresses active motion toward the minus end more effectively than movement toward the plus end; see the illustration in Fig. 2C and the corresponding force-velocity relations (Fig. 2C, lower panel). Note that an asymmetric impact of drag on active forward and backward stepping very similar to that suggested here was recently observed experimentally for the kinesin-14 Kar3 of *S. cerevisiae* (14). Importantly, drag has different effects on the Brownian and the active components of the model. While it symmetrically reduces Brownian motion without introducing any bias towards either end of the MT, the asymmetric drag response of active motion leads, for large enough values, to a reversal of the existing bias at zero force (towards the minus end). Moreover, the force-velocity curves for plus- and minus-end-directed motions intersect close to the stall force (Fig. 2C, lower panel). As we have inferred from the experimental data that the attractive forces are in the range of the stall forces of plus-end- and minus-end-directed motion of Cin8 as measured previously (9), an asymmetric drag response is likely to play a significant role in collective Cin8 dynamics, and could potentially lead to a switching of directionality in Cin8 clusters.

To test these ideas, we performed extensive stochastic simulations of the full model while accounting for all of the above features (see section B1 of the Supporting Information for a detailed description of the computational model and the choice of parameters). Strikingly, we observed that our model for Cin8 motion indeed exhibits directionality switching of Cin8 clusters. The kymographs obtained from our stochastic simulations clearly indicate a tendency for clusters to move towards the plus end of MTs (Fig. 2D), in remarkable agreement with experimental observations (Fig. 1B). In order to quantify this visual impression, we analyzed the stochastic trajectories obtained from the numerical simulation to calculate the mean displacement velocity and the diffusion coefficient (which are listed in Table SB3 in the Supporting Information). In agreement with experiments, single Cin8 molecules show a minus-end-directed mean displacement velocity, opposite to that of pairs and clusters. Importantly, the model quantitatively reproduces the measured displacement velocities, as well as diffusion coefficients, of single Cin8 motors, pairs and higher oligomers of motors to a very high degree of accuracy (Tables 1 and SB3).

Since our computational model faithfully captures directionality switching as characterized by the displacement analysis, we performed a more detailed analysis of the distribution of cluster sizes and motion. To this end, we determined the full statistics of cluster-size-(cluster intensity)-dependent motility in simulations and experiments by computing and measuring the average velocities and degree of cluster intensity for each trajectory of single Cin8 motors or clusters. Figure 3A shows that the resulting distribution obtained from simulations is in excellent agreement with the data obtained from experiments (Fig. 3B). Additionally, we found good agreement between experimental and simulation results for the clustering behavior at different Cin8 concentrations. Thus, increasing the Cin8 concentration *in silico*, as well as *in vitro*, led to an increased number of clusters (bright traces in kymographs) that slowly moved either in an undirected or plus-end-directed manner (Fig. 3C and D, top panels). This agreement is supported by the statistics of simulated and experimentally determined cluster intensity. Our model for collective Cin8 motion, which accounts for active as well as Brownian motion, produces concentration-dependent cluster-intensity distributions that agree with experimental data (Fig. 3C and D, bottom panels). The similarity of these results to those obtained with the model that accounts for Brownian motion only (cf. Fig 3C, and D and Fig. SB7) further supports the inference that clustering is only affected to a minor degree by active motion, as argued previously. Secondly, in our simulations we found a sharp transition between a regime with unstable clusters and one in which cluster formation is strongly favored with increasing Cin8 concentration (Fig. SB10). The concentration at which this sudden transition occurs agreed excellently with that which marks the onset of cluster formation in our experiments (see section B8 of the Supporting Information for additional data and the detailed analysis).

**Figure 3:**
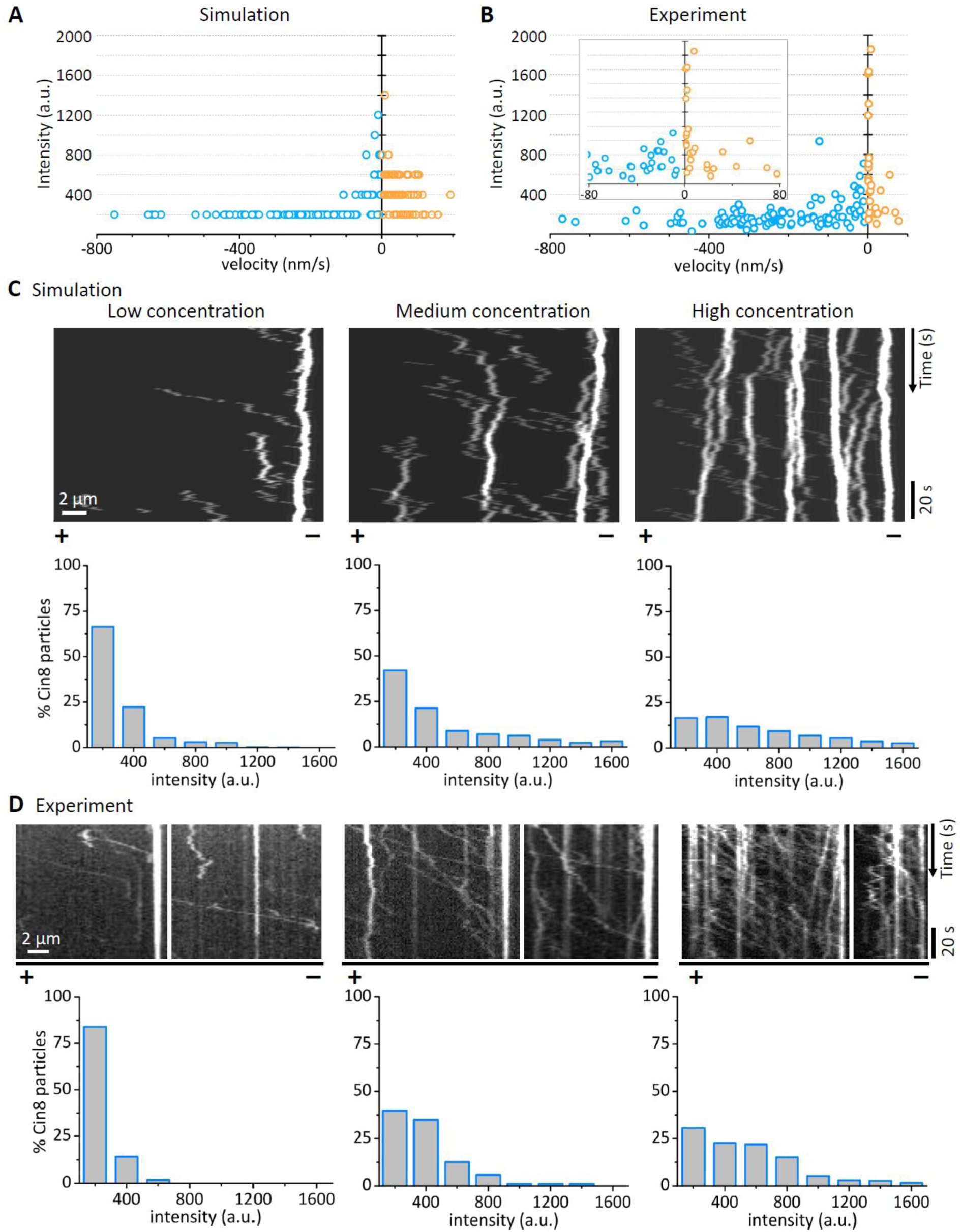
Comparison of simulation results with experimental data. **(A)** Relationship between velocity and intensity of Cin8-GFP motors obtained by simulations based on the computational model. Intensity is determined by multiplying the number of single Cin8 molecules in a cluster by 200 a.u., the maximal intensity of a single Cin8 tetramer, based on the photobleaching curves (Fig. 1A). **(B)** Same as **(A)**, but measured experimentally. Total n = 134; minus-end-directed n = 107, plus-end-directed n = 27. ⁓20% of the total motors are plus-end-directed, 70% of the total minus-end-directed motors are monomers. The majority (⁓88%) of plus-end-directed motors are clusters comprising >2 Cin8 molecules. Inset: Close-up of slow-moving Cin8 motors. In **(A)** and **(B)** blue and yellow circles represent (−) and (+) - end directed Cin8 molecules, respectively. **(C)** Concentration-dependent clustering in simulations. Top: Model-generated histograms of intensities of Cin8 motors bound to MTs at increasing concentration of Cin8. Bottom: Corresponding kymographs for different concentrations of Cin8, generated by theoretical simulations. **(D)** Experimental data for concentration-dependent clustering of Cin8 on single MTs. Top: Histograms of intensities of Cin8 motors bound to MTs at increasing concentration of Cin8. Bottom: Representative kymographs obtained from time-lapse measurements of Cin8 motility on MTs at the different Cin8 concentrations. Attachment rates in **(C)** (listed from left to right) were {3.62 · 10^−5^, 4.82 · 10^−5^, 9.04 · 10^−5^}, as fixed by the attachment rates measured experimentally at medium Cin8 concentrations (1-2 pM). A detailed description and a complete list of the parameters employed in **(A)** and **(C)** is found in section B2 and Table SB1 of the Supporting Information for. A description of the convolution method used to generate simulated kymographs is given in section B9 of the Supporting Information.

### A caterpillar mechanism for directionality switching of motion in Cin8 clusters

What is the basic mechanism underlying the observed directionality switching of Cin8 clusters relative to the behavior of single Cin8 motors? Key evidence emerged from additional simulations in which we suppressed Brownian motion of Cin8. Specifically, we found that directionality switching of clusters was absent when motors exhibited negligible Brownian motion (Fig. SB9), which points to a decisive role of Brownian motion in the directionality switching of Cin8 clusters. How the combined effects of diffusive motion and an asymmetric response of *active* motion to drag lead to clustering-induced directionality switching can be understood by the following mechanism, illustrated in Fig. 4.

**Figure 4:**
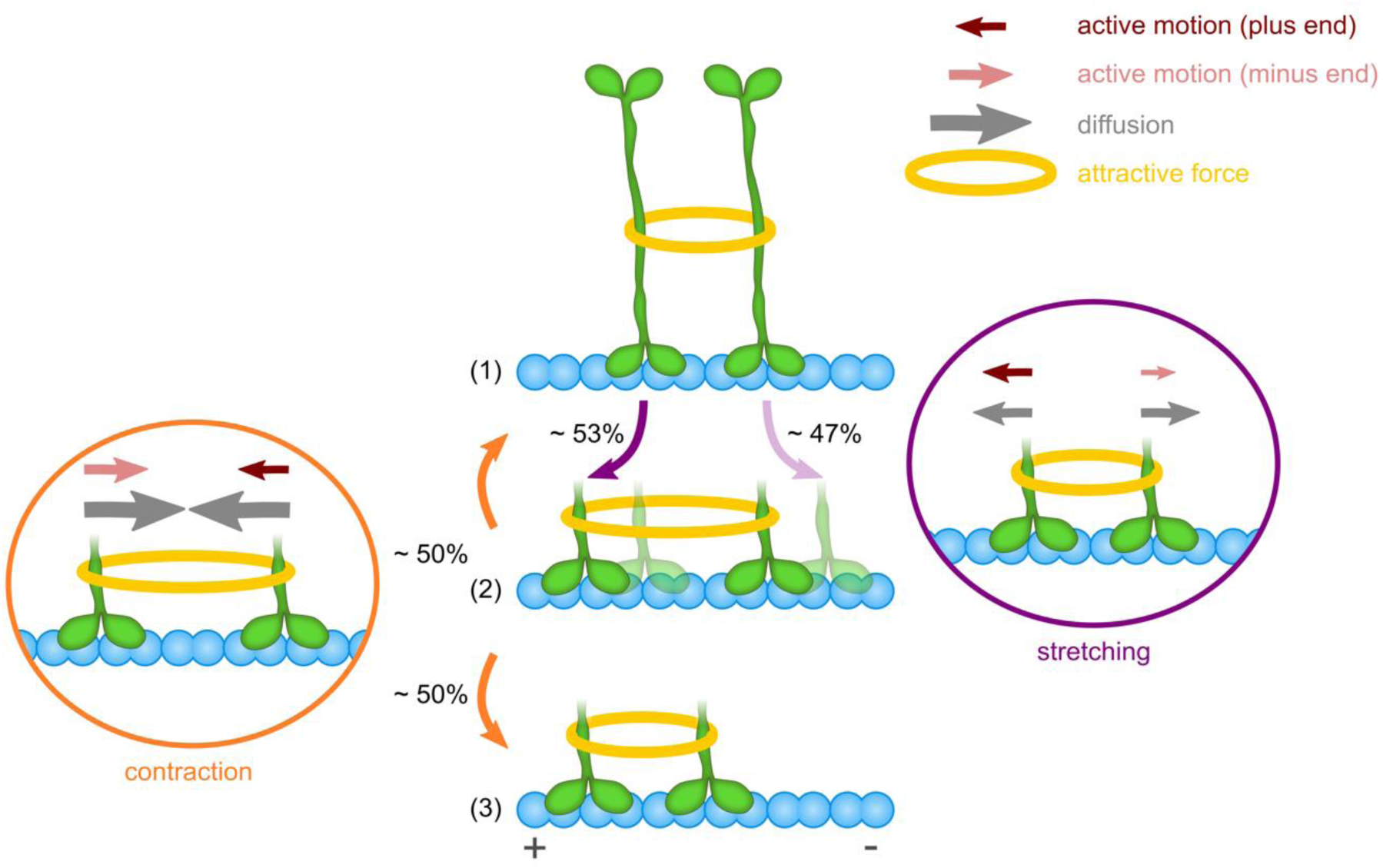
Model for plus-end directional preference of Cin8 clusters. A fully contracted cluster of two Cin8 motors (1) may expand by movement of either motor away from the other (cluster stretching, right panel). Stretching is constrained by the attractive interaction (yellow ring), which opposes both diffusion (gray arrows) and active motion (red/pink arrows). Conversely, motor clustering implies that diffusion is suppressed exponentially and active motion is biased towards the plus end owing to the assumed asymmetric response of the active motion of Cin8 to opposing forces (Fig. 2C). Consequently, stretching is slightly more likely (53%) to occur due to plus-end-directed motion of the left motor [opaque configuration in (2)] than by minus-end-directed motion of the right motor [transparent configuration in (2)]. Once the motors have moved further apart (2), the pair most likely contracts again [process (2)→(1) or (2)→(3)]. In the case of contraction (left panel), the attractive force supports both active motion and diffusion. Hence, active motion now shows a minus-end-directed bias (see Fig. 2C for supporting illustration). However, diffusive motion dominates cluster stretching, as it is supported (and not suppressed) by the attractive interaction. Since diffusive motion is inherently undirected, cluster contraction occurs almost equally often by motion of either motor (50.5% left motor; 49.5% right motor). As cluster stretching shows a weak bias towards the plus-end while cluster contraction is almost unbiased, a pair of motors is more likely to move in a caterpillar-like fashion toward the plus end [sequence of processes (1)→(2)→(3)] than the minus end. The sizes of the arrows differ for illustrative purposes only; they are not proportional to the model parameters (see section B6 of the Supporting Information for a detailed description of stochastic process and a computation of the conditional probabilities).

We start from a configuration in which two interacting Cin8 molecules are immediately adjacent to each other on the MT (Fig. 4, part (1)). The center-of-mass of this cluster can change if either moves away from the other, which we refer to as stretching (Fig. 4, right panel). Therefore, stretching can occur by either plus-end-directed motion of the left motor or minus-end-directed motion of the right motor. Both of these (stochastic hopping) processes are counteracted by the attractive interaction (yellow rings in Fig. 4) between the motors. The diffusive component of this motion (gray arrows) is suppressed by a Boltzmann factor in the same way for right and left motors. Due to the assumed asymmetry in the response of active motor motion to drag (Fig. 2C), the attractive interaction between the motors suppresses minus-end-directed active motion (pink arrows in Fig. 4) more strongly than plus-end-directed active motion. In other words, the left motor is more likely to move towards the plus end than the right motor is to shift in the direction of the minus end, thereby creating an overall bias for cluster motion towards the plus end. This process leads to a configuration in which the two motors have moved further apart (Fig. 4, part (2)). Now, they can not only diverge further from each other but move closer together, causing ‘cluster contraction’ (Fig. 4, left panel). Since the force between the motors is attractive, both diffusive and active motions are much more likely to draw them closer to each other than drive them further apart, so that one can effectively disregard the latter possibility. Importantly, for cluster contraction, the direction of motion of the motors corresponds to the direction of the forces at work, i.e. a pulling force instead of a drag then acts on each of the two motors. In that case, the active component of motion has the same bias as for zero load, i.e., the left motor is more likely to move towards the minus end than the motor on the right is to step toward the plus end (Fig. 2C, lower panel). Hence, this type of contractile motion creates a bias towards the minus end. Overall, the force-induced bias in the active motion acts in opposite directions during cluster stretching and cluster contraction, respectively.

These two opposing directional biases, however, do no cancel out, owing to the key role played by diffusive motion. According to the Arrhenius law, thermal Brownian motion is exponentially reduced when movement ensues against a force as compared to moving along the direction of a force with the same magnitude. As a consequence, this thermal component of motor motion is much more pronounced for cluster contraction than for cluster stretching (compare gray arrows in the left and right panels of Fig. 4). Moreover, Brownian motion is inherently undirected and therefore affects both motors in the cluster in the same (albeit mirror-symmetric) way. Adding up the active and thermal components of motion, this implies that Brownian motion has very different effects on the relative bias for cluster motion toward the plus end and minus ends. For cluster contraction, thermal motion is the dominating mode of motion, and therefore strongly reduces the relative bias in cluster motion towards the minus end caused by active motion. Consequently, contracting clusters are almost equally likely to move toward either end (50.5% motion of the left motor; 49.5% motion of the right motor). In contrast, during cluster stretching, thermal motion is exponentially suppressed, such that a clear bias for cluster motion towards the plus end ensues (53% motion of the left motor; 47% motion of the right motor); see section B6 in the Supporting Information for the computation of the conditional probabilities. Thus plus-end-directed motion of Cin8 clusters occurs because they preferentially stretch towards the plus end, while there is no pronounced directional bias during cluster contraction. Whenever plus-end directed cluster stretching is followed by plus-end directed cluster contraction, a wave-like motion of the motors towards the plus end occurs, reminiscent of *caterpillar motion* (sequence of processes (1)->(2)->(3) of Fig. 4).^2^ Importantly, the same arguments as made here for a pair of motors can be made for the two outermost motors of a cluster with more than two motors, such that these motors bias cluster motion towards the plus end. Unlike the case for a pair of Cin8, however, motility is further reduced by the motion of the motor(s) in the center of the cluster. As these motors experience forces from both directions, which cancel each other out, they behave like single motors with a large diffusive component in motion and a minus-end-directed bias. Thus, in agreement with experimental data (Fig. 1D and E, and Table 1), the motion of large clusters in the plus-end direction is slower than that of pairs of Cin8 (Table SB3).

### Asymmetric response to drag is supported by Cin8 motility experiments at low ionic strength

The analysis based on the theoretical model suggests that the asymmetric response of Cin8 to drag is the mechanism underlying directionality switching from minus- to plus-end-directed motility. While this conclusion follows indirectly from the comparison of the single-molecule motility experiments with simulations, we asked whether such an asymmetric response could also be observed directly. Single Cin8 motors are likely to experience increased drag while moving along a MT if their affinity for the MTs is increased. Such a change of affinity can typically be achieved by reducing the ionic strength of the buffer, since electrostatic interactions are less shielded under such conditions. Accordingly, we performed experiments at a salt concentration of 110 *mM* KCl. We indeed observed that decreasing the salt concentration drastically increased the number of MT-bound Cin8 motors compared to high-salt conditions at similar concentrations of Cin8 (Fig. 5A) and increased the landing rate of Cin8 (section SA2.3). This indicates that lowering the salt concentration increases the affinity of Cin8 for MTs, and hence results in higher drag on moving Cin8 motors. To compensate for the increased affinity, experiments under low salt conditions were performed with ⁓5-fold lower Cin8 concentration (Fig. 5B-E).

**Figure 5:**
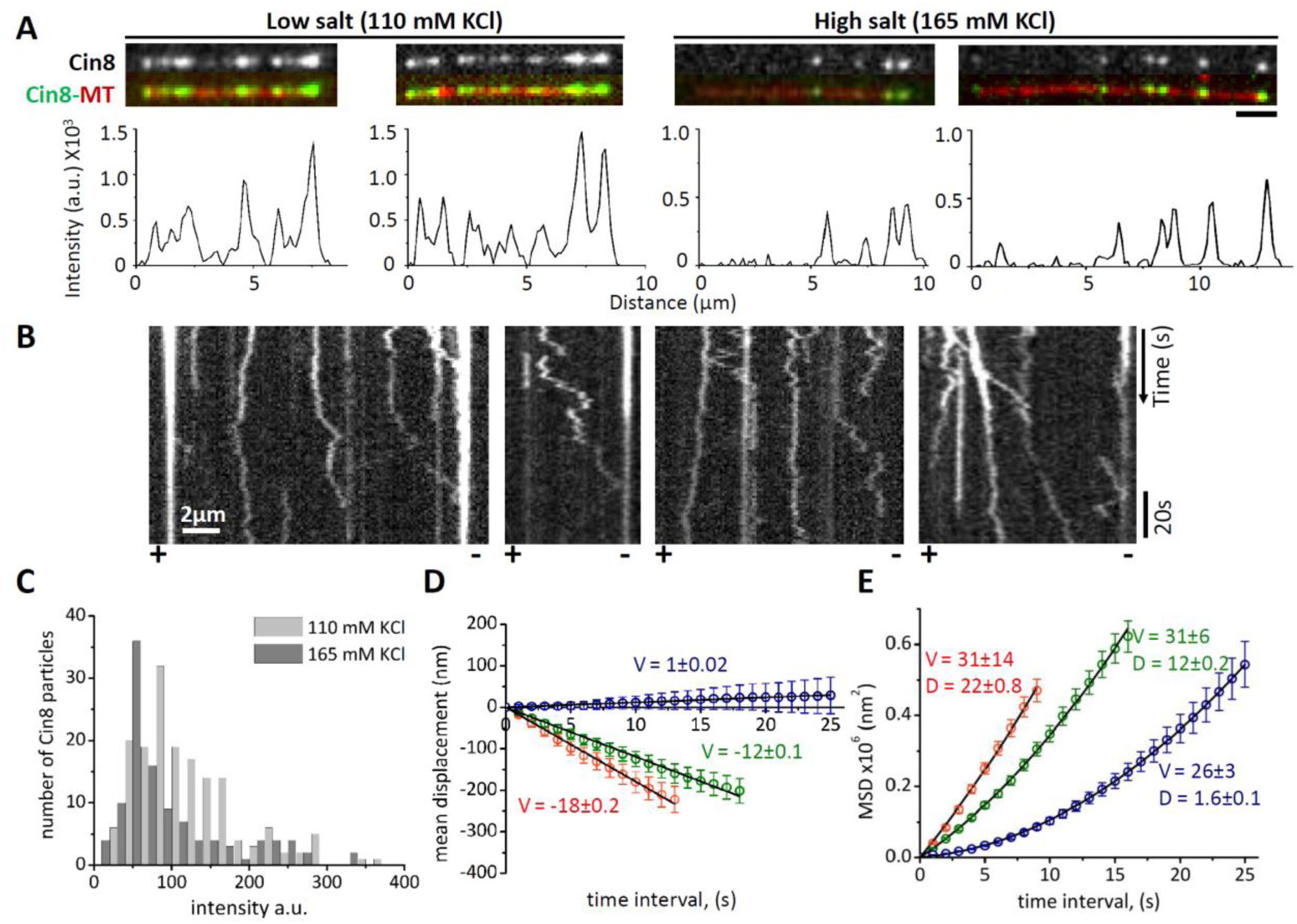
Motility of Cin8 under low-salt conditions. **(A)** MT-bound Cin8 in the presence of low or high salt concentration. **Top:** Representative images of MT-bound Cin8 at salt concentrations indicated at the top. **Bottom:** corresponding Cin8-GFP intensity profile of MT-bound Cin8 motors. Cin8 concentration is 1.3 pM and 2 pM for the low and high salt experiments, respectively. **(B)** Representative kymographs of Cin8-GFP motility along single polarity-marked MTs (polarity is indicated at the bottom) in the presence of 110 mM KCl and 1 mM ATP. **(C)** Intensity distribution histogram of MT-bound Cin8 at high (165 mM KCl, dark gray) and low salt (110 mM KCl, light gray). **(D and E)** Mean displacement (MD) **(D)** and mean square displacement (MSD) **(E)** for single Cin8 molecules (red), pairs of Cin8 (green) and higher oligomers (blue). The black solid lines are linear (MD = vt) (MD) or statistically weighted second-order polynomial fits (MSD = v^2^t^2^+2Dt) (MSD). Values of the mean velocity (V, nm/s) and the diffusion coefficient (**D**, nm^2^/s (x10^3^)) are indicated. Error bars represent SEM. **(B-E)** Cin8 concentrations are 2 pM and 0.4 pM for high and low salt, respectively.

In agreement with previous reports (7,8,29,30), we found that decreasing the salt concentration increases the fraction of plus-end-directed Cin8 trajectories and considerably diminishes the average displacement velocities (Fig. 5B, D, SA2b and Table 1). However, it had not been previously established whether this effect is caused by stronger Cin8-MT interaction or Cin8 clustering induced by low ionic strength. Our ability to analyze the motility of single molecules, pairs and oligomers of Cin8 separately enabled us to differentiate between these different effects on Cin8 motility. Importantly, in agreement with our hypothesis of an anisotropic response of active motion to drag, our analysis revealed that it is not clustering per se, but an increased drag that switches the directionality of Cin8. Under high-salt conditions only 3% of the trajectories of single Cin8 molecules were plus-end directed (Fig. SA2a), while 33% of single Cin8 trajectories showed switched directionality at low ionic strengths (Fig. SA2b). In addition, we were able to estimate the additional drag experienced by a single Cin8 motor moving on the MT in low-salt buffer. According to the Arrhenius law, drag suppresses diffusive motion by an exponential factor of *e*^−Δ*E*/*k*^*_B_^T^*, where Δ*E* is the additional energy barrier a motor has to overcome when moving from one binding site on the MT to the next. We estimated that the energy barrier must be surmounted approximately halfway between two tubulin dimers, i.e. *⁓*4 *nm*. Then, following the Arrhenius law, the 5-fold decrease in the diffusion coefficient observed for single Cin8 molecules in low salt relative to high salt (Fig. 5E and Table 1) implies an additional drag of approximately 1.6 *pN*. This value agrees with our modeling hypothesis that forces of this magnitude which antagonize motion switch the directionality of Cin8 (see Fig. 2C, lower panel).

Taken together, we found that consistent with our model, trajectories of single Cin8 motors become increasingly biased towards the plus-end direction along with experiencing a larger drag at low ionic strengths. Therefore, not motor clustering per se but drag or any external force that opposes motion can switch the directionality of Cin8.

## Discussion

Although bidirectional motility of kinesin motors was discovered nearly a decade ago (6,7), the underlying mechanisms had remained obscure. Our analysis shows that clustering-induced switching of motor directionality arises from an interplay between active motion in the minus- and plus-end direction, a significant diffusive component, and an asymmetric response of active motion to forces that oppose this motion. This implies that unique molecular features of the motor enable stepping in two directions. Specifically, the kinesin neck-linker has been shown to be the major structural determinant that differentiates between the directionality of the N-terminal (plus-end-directed) and the C-terminal (minus-end-directed) kinesins (31-35) and modulates the kinesin stepping mechanism and processivity (36,37). Hence, to enable stepping in two directions, one might speculate that the neck-linker of Cin8 exhibits a higher degree of flexibility than those of unidirectional kinesin motors. In addition, it has recently been demonstrated that the large insert in loop 8 within the catalytic domain of Cin8, which is important for directionality switching (7), is required for the non-canonical binding of 3-4 motor domains of Cin8 per tubulin dimer, probably via an attractive motor-motor interaction (38). In light of this finding, we hypothesize that loop 8 of Cin8 may be one of the factors responsible for the weak attractive forces between Cin8 tetramers that enable them to form clusters. In addition, the large loop 8 of Cin8 can interact with MTs, which may well affect the level of drag experienced by motors as they move along a MT and could therefore contribute directly to directionality switching. Finally, it has been shown that deletion of the C-terminal tail of Cin8 induces slow and bidirectional motility under high ionic strength conditions (29), similar to the motility of clusters reported here. Thus, we propose that the C-terminal tail of Cin8 may be one of the intramolecular factors that regulate clustering of Cin8 and thus favor its capacity for bidirectional motion.

The asymmetric force-velocity relation of bidirectional motion of Cin8 proposed in this study explains why directionality switching can be triggered by a multitude of different factors, as observed previously. The asymmetric response to drag explains directionality switching due to clustering of motors as well as changes in motor concentration and/or ionic strength. Importantly, it readily explains several seemingly unrelated observations previously reported for directionality switching (5-7,9,10,12). First, drag acting on motors can originate not only from interactions between Cin8 motors on MTs and their interactions with MTs, but also in multi-motor gliding and antiparallel sliding assays, in which a large ensemble of motors interact with the same MT. In such assays, changing the surface density of molecular motors alters the binding strength of the moving MT with the motor-covered surface. This in turn is expected to affect the drag underlying the gliding motion. Hence, an asymmetric response of active Cin8 motion to drag also explains the directionality switching in surface-gliding and antiparallel-sliding assays reported for the two kinesin-5 motors Cin8 and Kip1 (6,7,10). These conclusions are further supported by recent simulations of a gliding assay (39) in which a directionality switch in the motion of gliding MTs was observed when the displacement velocity of Cin8 motors depended asymmetrically on opposing forces – which is equivalent to our basic assumption of an asymmetric response of active motion to drag. Moreover, drag acting on a motor can also result from interactions with other proteins (with or without motor activity) that are bound to MTs. Our analysis suggests that such interactions lead to directionality switching, in agreement with recent findings indicating that crowding on MTs with non-motor proteins switches the directionality of the bidirectional *S. pombe* kinesin-5 Cut7 (12). Finally, directionality switching has also been observed for mitotic kinesin-14 motors (13,14), and an asymmetric response to force has been directly demonstrated for the *S. cerevisiae* kinesin-14 Kar3 (14). Thus, the suggested mechanism of an asymmetric response of motor motion to drag may also explain directionality switching of other kinesins, therefore providing a unified view of directionality switching of kinesin motors.

## Materials and Methods

The MT polymerization and *in vitro* single-molecule motility experiments were performed as previously described (10). Cin8 cluster size determination is achieved by monitoring the photobleaching of GFP attached to Cin8. Cin8 motility is characterized using mean displacement (MD) and mean squared displacement (MSD) analysis. The details are described in the Supporting Information.

## Data Availability

All relevant data are presented in this paper and the accompanying Supporting Information files.

## Acknowledgements

This research was supported in part by the Israel Science Foundation (ISF-386/18) awarded to L.G.; the National Science Foundation (NSF-1615991) and United States - Israel Binational Science Foundation grant (BSF-2015851), awarded to L.G. and J.AB.; the National Institutes of Health (grant NIH-R01-GM11283, awarded to J.A.B.; and the Deutsche Forschungsgemeinschaft (DFG, German Research Foundation) via project B02 awarded to E.F. within the Collaborative Research Center ‘Forces in Biomolecular Systems’ (SFB 863 – Project ID 111166240).

## Supporting information

### A Experimental

#### A1 Experimental Materials and Methods

##### A1.1 Overexpression and purification of Cin8-GFP

Cin8-GFP was overexpressed and purified from a protease-deficient *Saccharomyces cerevisiae* strain containing a 2-μm plasmid for Cin8-GFP–6His, expressed under the *pGAL1* promoter [LGY 4093 (*MATα, leu2-3,112, reg-1-501, ura3-52, pep4-3, prb1-1122, gal1*, (*pOS7: 2μ, LEU2, PGAL1-CIN8-GFP-6HIS*))] as previously described (1, 2). Briefly, *S. cerevisiae* cultures were grown in a minimal medium supplemented with 2% raffinose; Cin8 expression was induced by the addition of 2% galactose. Cells were pelleted and then ground in liquid nitrogen. The ground cells were thawed and centrifuged, and the supernatant was loaded onto a Ni-NTA column pre-equilibrated with equilibration buffer (50 mM Tris-HCl, 30 mM Pipes, 500 mM KCl, 10% glycerol, 1.5 mM β-mercaptoethanol, 1 mM MgCl2, 0.1 mM ATP, 0.1% Triton X-100, pH 8). After washing (with equilibration buffer supplemented with 25 mM imidazole), Cin8-GFP-6His was eluted with elution buffer (50 mM Tris-HCl, 30 mM Pipes, 500 mM KCl, 350 mM imidazole, 10% glycerol, 1.5 mM β-mercaptoethanol, 1 mM MgCl2, 0.1 mM ATP, 0.1% Triton X-100, pH 7.2), and the eluted fractions were purified by size-exclusion chromatography using a Superose-6 (10/300) column (GE Healthcare). The fractions corresponding to the Cin8-GFP tetramer were collected and analyzed by SDS-PAGE and western blotting for detection of Cin8-GFP. The selected fractions were aliquoted, snap frozen, and stored until use at −80°C.

##### A1.2 *In-vitro* single molecule Cin8 motor motility and photobleaching

Assays for microtubule single-molecule motor motility were performed as previously described (1). Briefly, GMPCPP (guanylyl 5’-α,β,-methylenediphosphonate)-stabilized, fluorescently labeled, biotinylated MTs were polymerized with 1 mg/ml tubulin, 0.08 mg/ml biotinylated tubulin and 0.08 mg/ml rhodamine-labeled tubulin, in the presence of 1 mM GMPCPP for 1 h at 37°C in general tubulin buffer (GTB; 80 mM Pipes, 0.5 mM EGTA, 2 mM MgCl2, pH 6.9). For plus-end labelling of MTs, an additional 0.08 mg/ml rhodamine-labeled tubulin was added after 1 h, and the mixture was incubated at 37°C for another 45 min (1). Commercially available tubulin (Cytoskeleton Inc., USA) was used.

Coverslips cleaned in piranha solution and subsequently treated with 0.1% dimethyldichlorosilane in trichloroethylene were assembled into flow cells of volume 10 µl by using double-sided tape. The salinized coverslip surface was coated with 1 mg/ml biotinylated bovine serum albumin (Sigma-Aldrich) and then treated with 1 mg/ml NeutraAvidin (Life Technologies) to attach biotinylated MTs. Following the attachment of pre-polymerized biotinylated MTs and washings, 20 µl of Cin8-GFP in motility buffer (50 mM Tris-HCl, 30 mM PIPES, 165/110 mM KCl, 2 mM MgCl2, 1 mM EGTA, 50 µg/ml casein, 1 mM DTT, ATP/ADP, 10 mM glucose, 100 µg/ml glucose oxidase, 80 µg/ml catalase, pH 7.2) was added into the flow cell. The final concentration of Cin8-GFP in the flow cell was 1-2 pM unless otherwise indicated. MTs and Cin8-GFP were imaged using a Zeiss Axiovert 200M inverted microscope equipped with an sCMOS camera (Andor). Motility data was acquired using MicroManager controller software (3). For Cin8-GFP motility and photobleaching experiments, 90-110 images were captured at 1 frame/second with an exposure time of 800 ms. Using 2×2 binning, the pixel resolution was 124 nm/pixel. Image analysis was performed using ImageJ-Fiji software (4). All measurements were acquired in at least three independent experiments using three different protein preparations for experiments at high KCl concentration and two different protein preparations for experiments at low KCl concentration.

##### A1.3 Experimental motility data analysis

Directionality of the MTs was assigned on the basis of plus-end labeling and/or the direction of fast-moving minus-end-directed Cin8 molecules (1). Kymographs were created using ImageJ-Fiji software. Mean displacement (MD) and mean squared displacement (MSD) analyses were performed as previously described (5-7). The coordinates of motile Cin8-GFP molecules and clusters were determined with the TrackMate plugin of the ImageJ-Fiji software or by manual tracking of the intensity center over time. Only those Cin8-GFP motors (“Cin8 motors” refers to moving Cin8 particles of unspecified cluster-size) that moved more than 3 pixels were considered motile, and only those with motility times > 4 s were tracked. Movements of the same molecule that included more than 6 s stall were considered as two separate movements. The number of Cin8-GFP motors traced in each size category is indicated in Table 1 of the main text. The coordinates obtained were used to calculate the Cin8-GFP displacement for each time interval. The MD and the MSD values were obtained by averaging the displacements and squared displacements, respectively, calculated for all motility recordings of Cin8-GFP on the MTs. Mean velocities (v) and diffusion constants (D) were derived by fitting the average MSD values to a weighted second-order polynomial equation MSD = v^2^t^2^ + 2Dt for facilitated directional motility conditions and to the linear equation MSD = 2Dt for purely diffusive motility conditions (7). Mean velocities (v) were also derived by fitting the MD functions to the linear equation MD = vt. For each time point, the average values and the standard error of the means (SEM) are indicated (Fig. 1D&E, main text). MSD data was plotted with statistically weighted 2nd-order polynomial MSD curves, by using the Origin (OriginLab) software, in such a way as to give higher weight to the initial time points, as these points represent more data. The MSD plot fitting for the motility of pairs of Cin8 molecules at 165 mM KCl and 1 mM ATP was performed after 15 s so as to exclude the initial nonlinear part of the curve. The v and D values for single Cin8 molecules, pairs of Cin8 molecules, and oligomers of Cin8 obtained from the MSD and MD analyses are summarized in Table 1 of the main text. Each motor was categorized as MT-plus-end directed if its net displacement was in the plus-end direction and if it remained continuously plus-end directed for at least three quarters of the length of its overall run. All other moving motors were classified as minus-end directed. The MSD values were also plotted versus time intervals on a log-log scale and fitted to a linear equation (Fig. SA6) to distinguish between diffusive and facilitated directional motility conditions. Movements with slopes >1.1 are considered as significant facilitated directional motility conditions (5, 6)

#### A2 Characterization of Cin8 motility and clustering

##### A2.1 Determination of Cin8 cluster size

To determine the size of Cin8 clusters, we first performed photobleaching experiments to determine the contribution of single GFP molecules to the total intensity of these clusters. We followed the fluorescence intensity of Cin8-GFP as a function of time within a circle of radius 4 pixels, using the TrackMate plugin of the ImageJ-Fiji software (8), following correction for uneven illumination and background subtraction. We observed that single photobleaching steps, probably representing photobleaching of 1 GFP, caused a reduction of ⁓50 a.u. in fluorescence (Fig. 1A, main text). Since each Cin8 tetramer contains four GFP molecules, all the Cin8 motors having an intensity ≤200 a.u. are likely to be single tetrameric Cin8 molecules.

The intensity distribution analysis of Cin8-GFP motors was consistent with the determination of the cluster size of Cin8-GFP from the photobleaching experiments. The intensities of all the fluorescent Cin8-GFP motors in the first frame of a time-lapse sequence were measured as described above. The major peak of the histogram of Cin8-GFP motor intensity distribution was fitted to a Gaussian curve (Fig. SA1). The center of the Gaussian peak lay at ⁓120 a.u., which was consistent with the average intensity of single Cin8 molecules containing one, two, three, or four fluorescent (non-bleached) GFP molecules, with each fluorescent GFP molecule contributing ⁓50 a.u. to the total intensity. Thus, the Cin8 population within this Gaussian peak, which constituted ⁓70% of the total Cin8 population, probably represents single Cin8 molecules. In addition, measures were taken to minimize the effect of GFP photobleaching on the determination of the Cin8 cluster size. First, we determined the lifetime of a GFP molecule before photobleaching under our experimental conditions as 26±4 (SEM) s (n=22). Consequently, based on this estimation, all our motility-cluster size measurements were performed only on those Cin8 motors that moved within the first 30 s of each measurement. Finally, for motile Cin8 molecules and clusters, we measured the fluorescence intensity only in the first frame of their appearance, thereby significantly reducing the likelihood of photobleaching. By this method, we assigned intensity ranges of Cin8 motor fluorescence as <200, 200-400 and >400 for single Cin8 molecules, pairs of Cin8 molecules, and Cin8 oligomers, respectively.

**Figure SA1.**
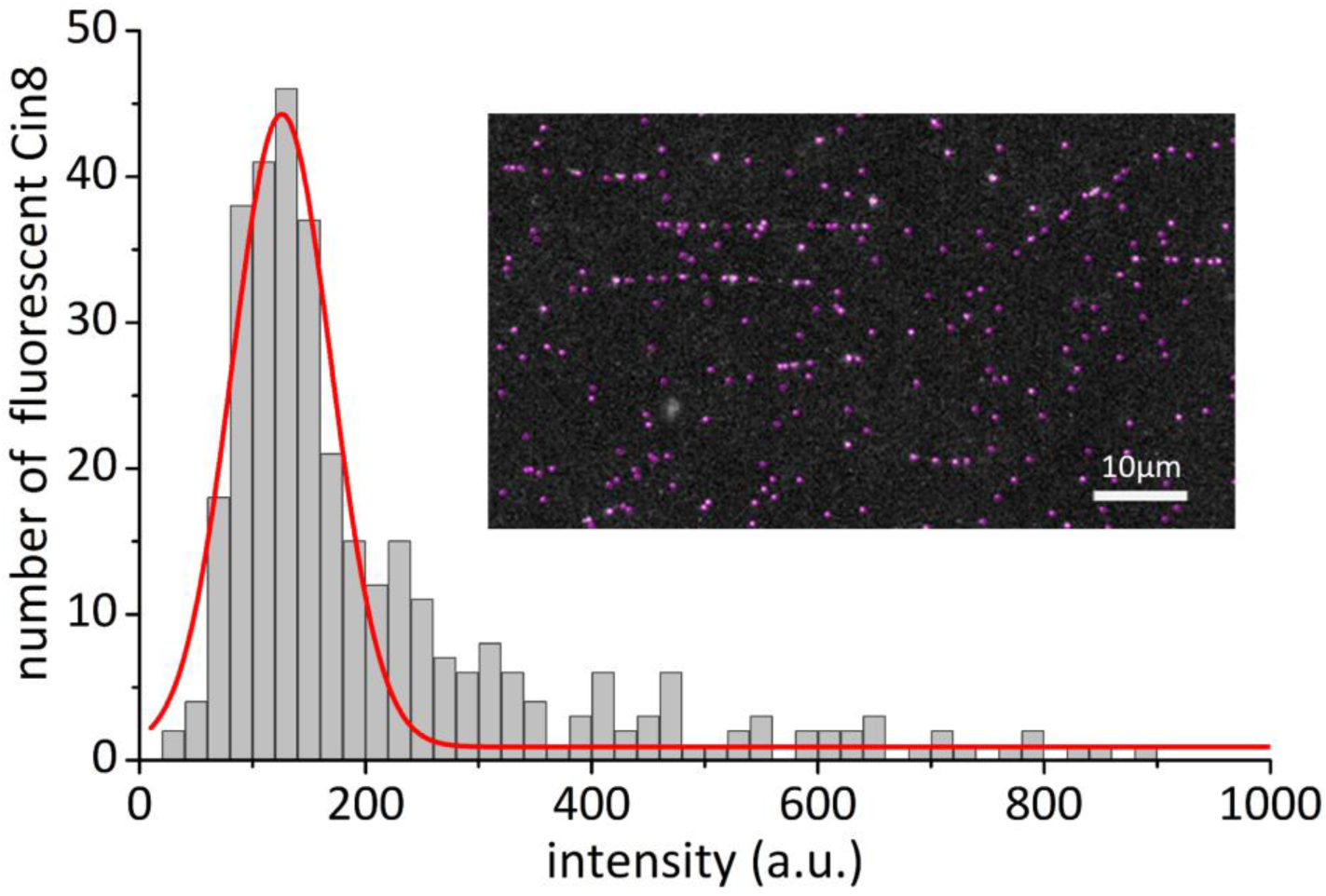
Intensity distribution of all Cin8 motors in the first frame of a time-lapse sequence (inset). The Gaussian peak (red) represents single Cin8 molecules constituting ⁓70% of the total Cin8 population. This peak is centered at ⁓120 a.u., which is the average intensity of a single Cin8 tetramer containing either 1, 2, 3 or 4 fluorescent GFP molecules, with each GFP molecule contributing ⁓50 a.u. to the total intensity. Accordingly, the maximal fluorescence of single Cin8 tetramers is 200 a.u.

For cluster size vs velocity analysis (Fig. 3B, main text), the velocity of each Cin8-GFP particle was determined by measuring the slope of each end-to-end motility event on the kymograph. The intensities of MT-bound Cin8-GFP motors were measured in the same manner as for the intensity distribution analysis described above, but only for the motile MT-attached Cin8 motors. The intensity profile of MT-bound Cin8-GFP motors was determined using line-scan analysis in ImageJ-Fiji software and plotted using the Origin (Originlab) software (Fig. 5A, main text).

##### A2.2 Characterization of cluster behavior

We found that while individual Cin8 tetramers predominantly move towards minus end of MTs, pairs and higher oligomers can switch directionality and move with greatly reduced velocity as compared to individual Cin8 molecules (Fig. SA2). Interestingly, we also observed that these MT-bound clusters of Cin8 move along MTs and sometimes split into two, and both products then move independently along the MT. Similarly, two Cin8 molecules can merge while moving on an MT to form a larger cluster that retains the characteristic motility of Cin8 (Fig. SA3).

In addition, we found that clustering is a concentration-dependent phenomenon. To estimate the effect of Cin8 concentration on clustering, we determined the intensity distribution (i.e., cluster size distribution) of the MT-bound Cin8 molecules in the presence of 1 mM ATP, 7.5 µM ATP or 1 mM ADP. We found that with increasing Cin8 concentrations, the intensity distribution for MT-bound Cin8 shifted towards higher-order oligomers (Fig. 3D, main text; Fig. SA4 and SA5), indicating that at higher concentrations, Cin8 has a high propensity to form larger clusters, irrespective of the presence of ATP or ADP, or the ATP concentration. With hardly any Cin8 clusters being found at the low Cin8 concentrations and large numbers of clusters being present at higher Cin8 concentrations, it is evident that clustering is a dynamic process with an equilibrium between single and clustered Cin8.

**Figure SA2.**
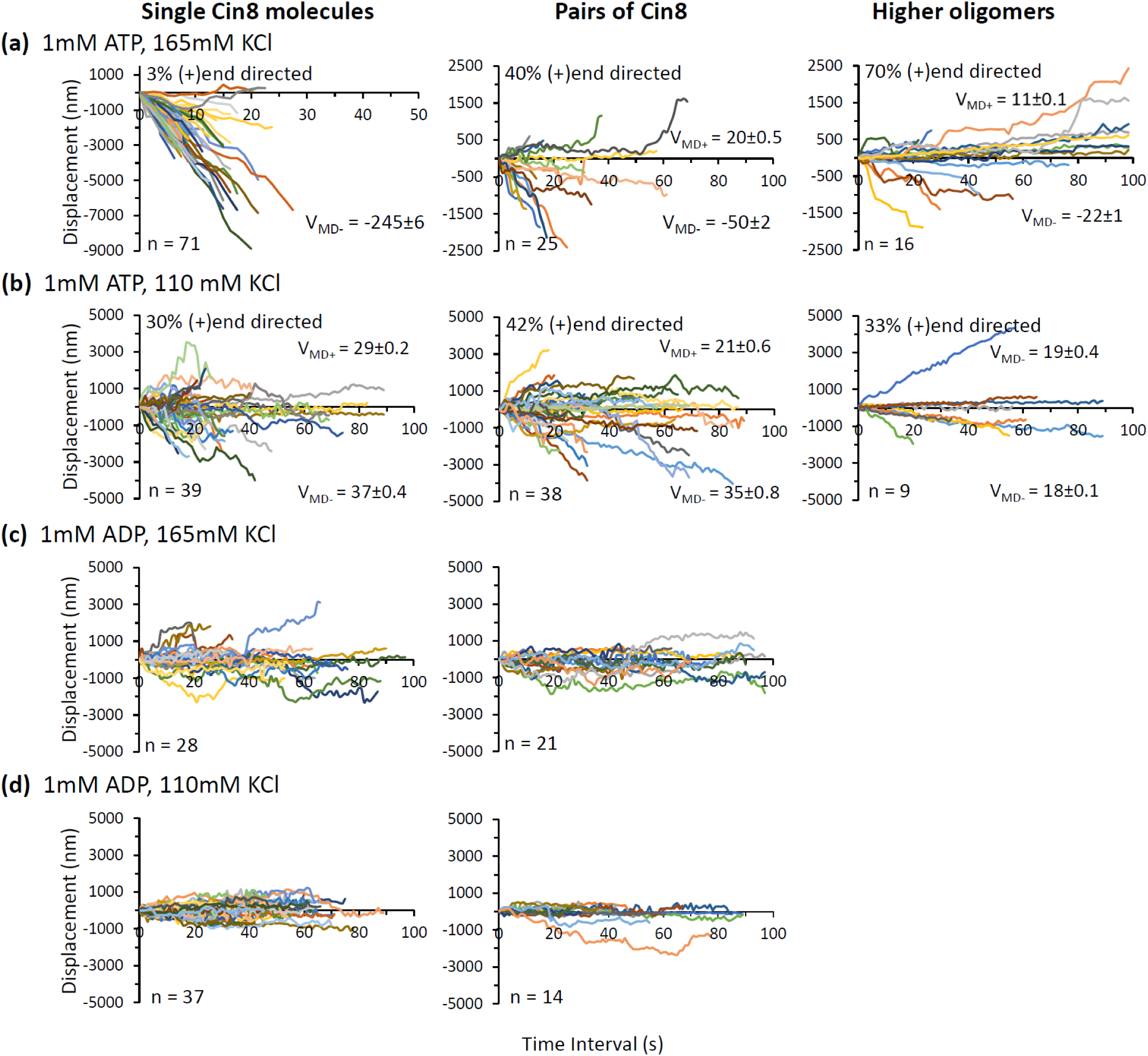
Displacement of Cin8-GFP motors. Displacement trajectories of Cin8 motors and clusters of different sizes (indicated at the top) in the presence of: **(a)** 1 mM ATP and 165 mM KCl (high salt); **(b)** 1 mM ATP and 110 mM KCl (low salt); **(c)** 1 mM ADP and 165 mM KCl (high salt); (**d)** 1 mM ADP and 110 mM KCl (low salt). n is the total number of Cin8 motors in each category. VMD+ and VMD-represent the velocity of (+) and (−) end-directed Cin8 motors in nm/s, respectively. The percentage of (+) end-directed Cin8 motors in the presence of ATP is indicated at the top left of each plot.

**Figure SA3.**
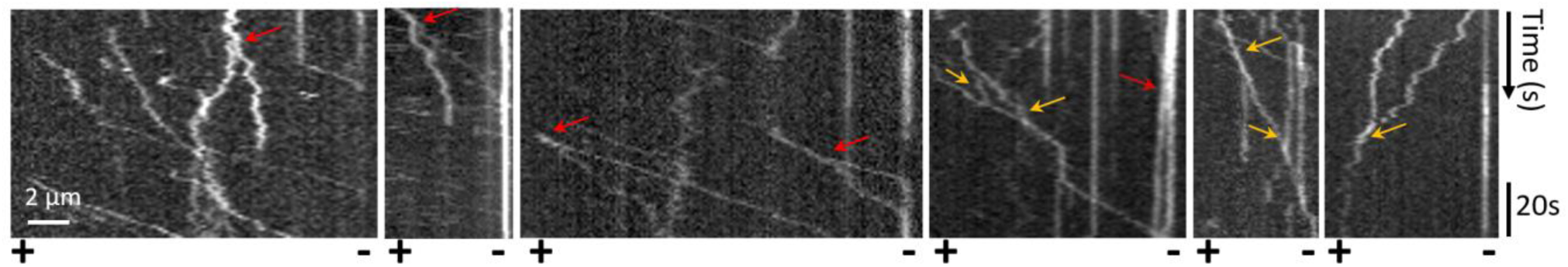
Dynamic clusters of Cin8-GFP. Reperesentative kymographs depicting dynamic clusters of Cin8-GFP, which split (marked with red arrows) into smaller clusters or individual Cin8 molecules, or merge (marked with yellow arrows) to form larger clusters while moving on single MTs.

**Figure SA4.**
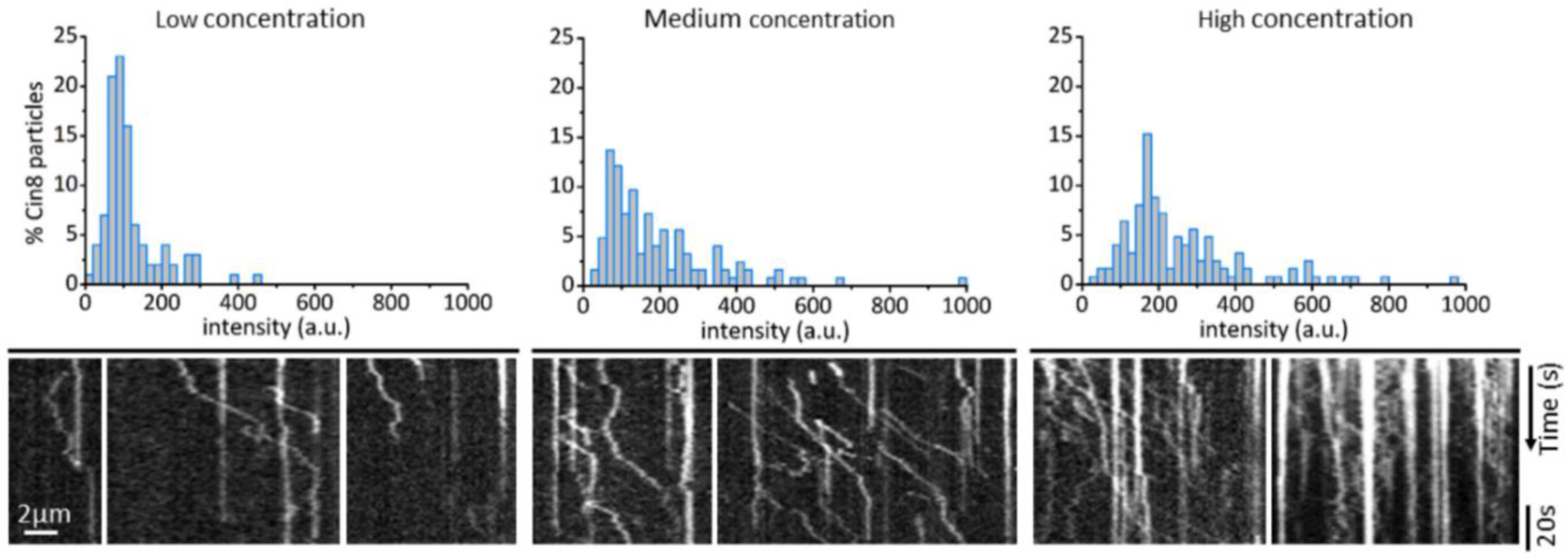
Concentration-dependent clustering of Cin8 on MTs in the presence of 7.5 µM ATP. Top: Intensity histograms of Cin8 motors bound to MTs at three different Cin8 concentrations. Bottom: Representative kymographs of time-lapse measurements of Cin8 motility on MTs at the corresponding different Cin8 concentrations.

**Figure SA5.**
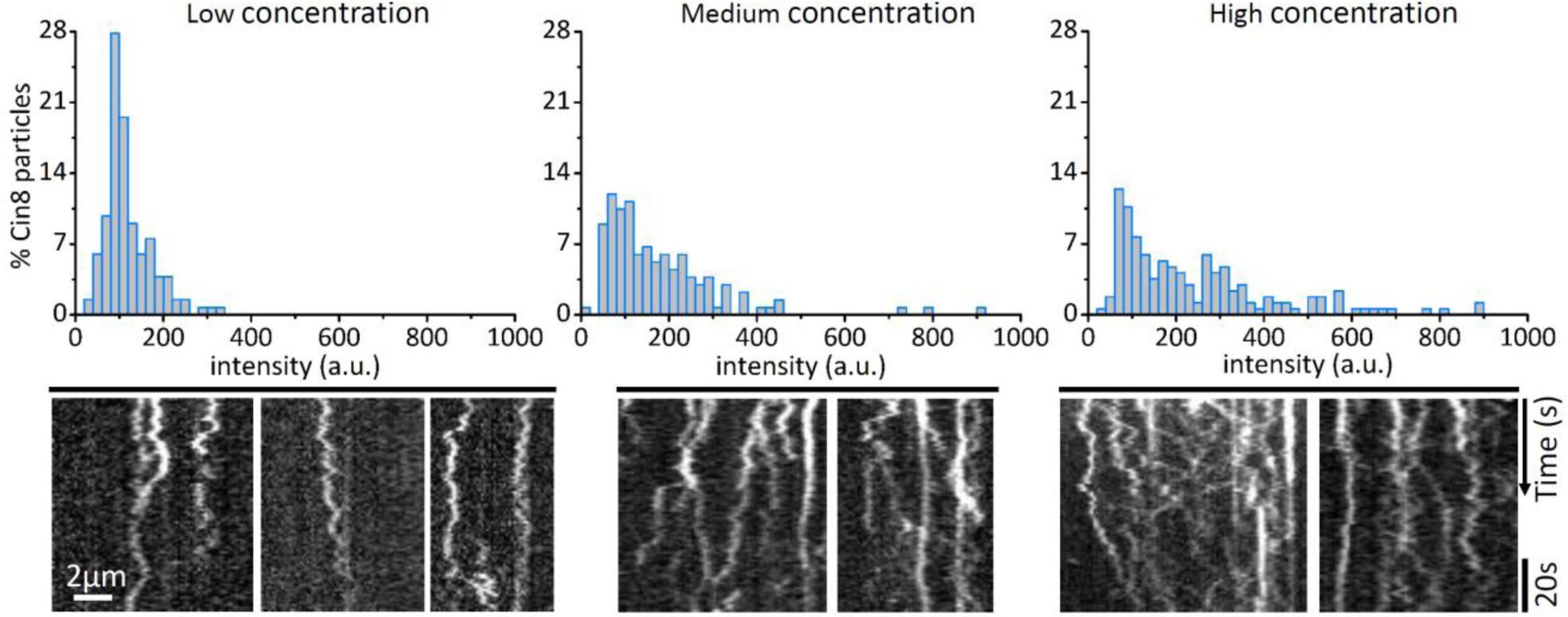
Concentration-dependent clustering of Cin8 on the MTs in the presence of 1 mM ADP. Top: Intensity histograms of Cin8 bound to MTs at three different Cin8 concentrations. Bottom: Representative kymographs of time-lapse measurements of Cin8 motility on MTs at the corresponding three different Cin8 concentrations.

**Figure SA6.**
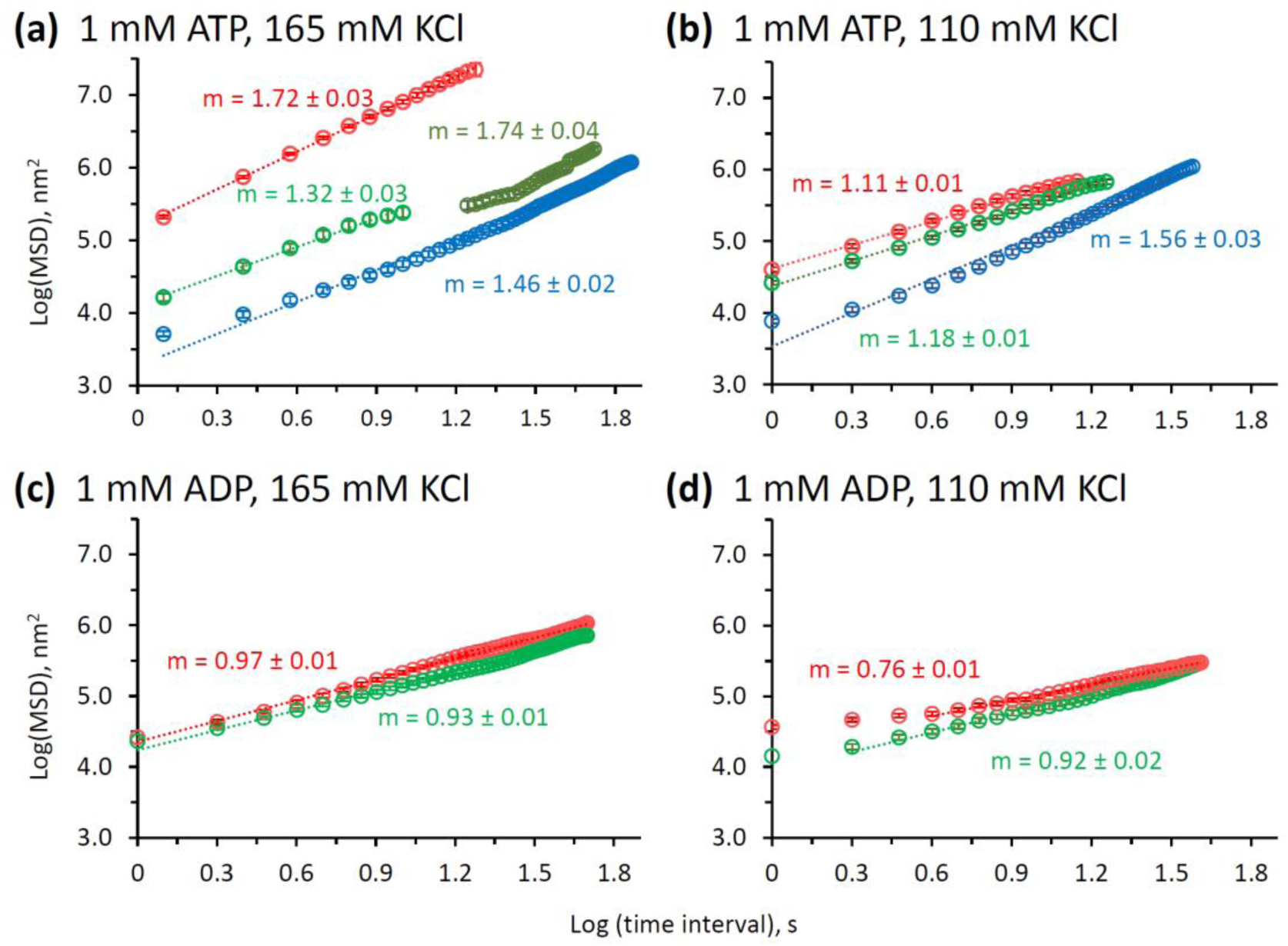
Log-log MSD analysis for Cin8 motility. Log (MSD) values were plotted vs. Log(time) for single Cin8 molecules (red), pairs of Cin8 (green), and higher oligomers (blue). The different salt and nucleotide conditions are indicated on top of each panel. Light green circles represent Cin8 dimers with short dwell times, and dark green circles represent Cin8 dimers with long dwell times. A slope of ⁓1 is consistent with purely diffusive motility (7). In the presence of ATP slopes >1 indicate a facilitated diffusive component. Dotted lines represent linear fits with the slope (m). Error bars represent SEM.

##### A2.3 Cin8 landing and attachment

The rates of Cin8 attachment to MTs were calculated as the total number of Cin8 motors attached per unit length of MT per unit time, whereas the landing rate was calculated as the number of Cin8 motors landing per unit length of MT per unit time after the first time frame of the experiment. Both attachment and landing rates were calculated for Cin8 motors that remained attached to MTs for at least 4 s. The attachment rate of Cin8 was 13.1±0.8 *10^−3^ and 27±3 *10^−3^ motors/(µm*s) for the high- and low-salt conditions, respectively. These findings indicate that lowering the salt concentration increased the affinity of Cin8 for the MTs and hence likely to increase the drag on Cin8 movement. Consequently, experiments under low-salt conditions were performed with a ⁓5-fold lower Cin8 concentration (Fig. 5, main text). Additionally, the Cin8 landing rate of 5.74 *10^−3^ motors/(µm*s) obtained in high salt condition experimentally was found to coincide with the particle landing rate at which clusters rapidly started to form in the simulations, namely, 5-6*10^−3^ motors/(µm*s) (Fig. SB10), indicating that the results of simulations were indeed consistent with those obtained experimentally.

##### A2.4 Dwell time

The dwell time was calculated as the time for which a motile Cin8-GFP particle remained attached to an MT (Table SA1). For the motors that remained attached to the MTs from the first time point recorded until the end of the acquisition, the dwell time was taken to be 90 s. The dwell time in the presence of ATP is dependent on Cin8 cluster size, with the pairs and higher oligomer clusters having ⁓4 times larger dwell time with respect to single Cin8 molecules, indicating an increased affinity of Cin8 clusters for MTs, as clusters have more MT binding sites with respect to single Cin8. Surprisingly, we found that in the presence of ADP Cin8 motors have large dwell times irrespective of whether they are present as single molecules or clusters, indicating an increased affinity of Cin8 for MTs in the presence of ADP.

**Table SA1.**
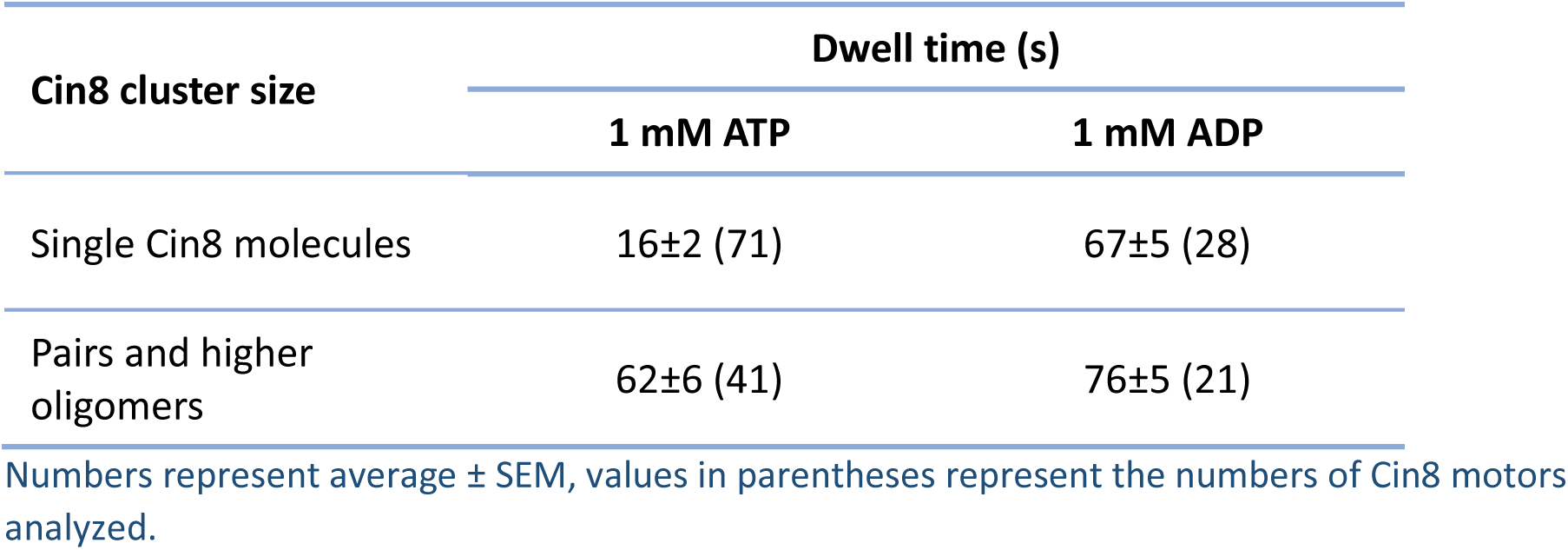
Dwell time (s) for moving Cin8-GFP motors at 165 mM KCl (high salt)

| **Cin8 cluster size** | **Dwell time (s)** |  |
| --- | --- | --- |
|  | **1 mM ATP** | **1 mM ADP** |
| Single Cin8 molecules | 16±2 (71) | 67±5 (28) |
| Pairs and higher oligomers | 62±6 (41) | 76±5 (21) |

Numbers represent average ± SEM, values in parentheses represent the numbers of Cin8 motors analyzed.

### B Theoretical model and simulations

#### B1 Description of the stochastic model for Cin8 dynamics

To model the collective dynamics of Cin8 motors on a microtubule (MT), we considered a one-dimensional lattice gas model (9). In the model, point-like *particles* that represent Cin8 motors populate a one-dimensional lattice that represents a MT. The lattice spacing *a* is chosen to reflect the size of a tubulin dimer, *a* = 8.4 nm (10). We assume that the stochastic dynamics of the system is described by a continuous-time Markov process. Each particle can show any of four basic stochastic processes, each characterized by a particular transition rate: (i) Plus-end-directed motion of a particle to the left neighboring lattice site, (ii) minus-end-directed motion of a particle to the right neighboring lattice site, (iii) attachment to a lattice site (i.e., a new particle is added to the lattice), or (iv) detachment from a lattice site (i.e., an existing particle is removed from the lattice). Each transition obeys an exclusion principle, which means that a particle may only be placed on or moved to a vacant lattice site. In this way, we account for steric interactions between Cin8 motors. In addition, particles are subject to an attractive interaction as soon as they come within an interaction range *R*. The transition rates of particles that are subject to interactions differ from those of non-interacting particles. Below, we detail the respective transition rates and explain how we modeled attractive interactions.

##### B1.1 Motion of non-interacting particles

Particles move stochastically to neighboring lattice sites in both the plus-end direction and the minus-end direction. Such movement may occur either by Brownian motion, described by a symmetric random walk, or by active, bidirectional motion, described by a biased random walk. The corresponding rates depend on whether a particle interacts with another particle, i.e., whether at least one other particle is located within a distance *R* from the first particle. For non-interacting particles (indicated by the subscript “0”) the respective rates are denoted by:

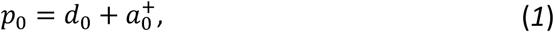

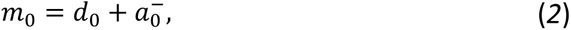

where *p*_0_ and *m*_0_ are the rates for movements in the plus-end and minus-end directions, respectively, *d*_0_ is the (symmetric) rate due to diffusion, and 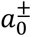 denote the rates for active plus- and minus-end-directed motion, respectively.

##### B1.2 Attachment and detachment of non-interacting particles

To account for interactions of an MT with the surrounding solution at a constant Cin8 concentration, we allow particles to bind and unbind to the lattice. Particles attach to vacant lattice sites at rate 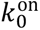. The rate of detachment of a non-interacting particle is denoted by 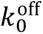.

##### B1.3 Attractive interactions

**Figure SB1.**
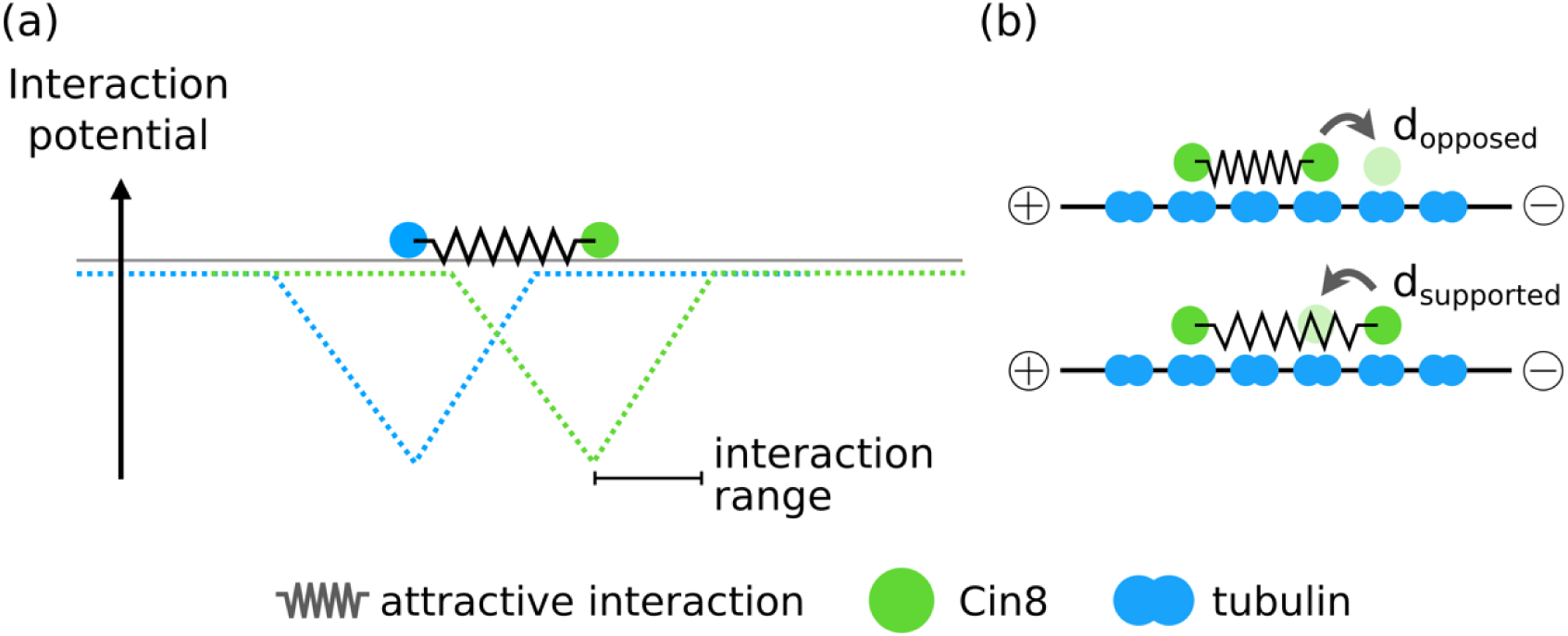
Illustration of particle interactions and their impact on the diffusive module of our model. **(a)** Interactions between particles were modeled as a constant force or, equivalently, as a linearly decreasing interaction potential (dashed lines) for distances shorter than a specific interaction range *R*. **(b)** For interacting particles, the attraction force, in general, affects the rates of movement to neighboring lattice sites. Because of detailed balance, the rate *d*^opposed^ for movement via Brownian motion against a force that arises from an interaction with another particle is related to the reverse rate *d^supported^* (motion via Brownian motion in the direction of the force) by the Boltzmann factor: *d*^opposed^/*d*^supported^ = exp(−*βΔE*) =: *δ*, where *ΔE* is the change in energy associated with moving the particles further apart over a distance *a* = 8.4 nm. Moreover, *β* = 1/k*T*, where *T* is the temperature and k is the Boltzmann constant. In the model, we implement only nearest-neighbor interactions. Note that forces are illustrated schematically by springs, although we assume constant forces for interacting particles.

In the model, two particles interact as soon as the distance between them becomes smaller than the interaction range *R.* As the specific functional relation between the interaction strength and the distance between two Cin8 motors is unknown but is expected to decrease monotonically with increasing distance, we assumed for the sake of simplicity that the interaction energy between two particles increases linearly for distances less than *R*, see Fig. SB1(a). This is equivalent to a constant central force *F*_int_ acting on both particles.^1^ Again, for the sake of simplicity, we assume that interactions are restricted to nearest neighbors. Hence, each particle can interact with at most two other particles. This assumption will also be justified *a posteriori*, as only small interaction ranges lead to biologically reasonable clustering behavior (see also the section “Model Parameters”). Due to our specific choice of particle interactions, a particle that interacts with two other particles, one to the left and one to the right, is not subject to a net force, as the interactions are in opposite directions and are therefore cancel out.

##### B1.4 Motion of non-interacting particles

Attractive forces between two particles favor motion that decreases the distance between them over motion that increases this distance. While we still assume that the overall rate of particle movement to neighboring lattice sites in the plus-end or minus-end direction is the sum of a rate due to passive diffusion and a rate due to active motion, attractive forces will, in general, affect the two components differently.

For Brownian motion of the particles, the rate *d*_supported_ at which either of two interacting particles moves the distance of a single lattice site closer to the other (supported by the interaction) is related to the rate *d*_opposed_ at which they move a single lattice site further apart (opposed by interactions, see Fig. SB1(b) for an illustration) via detailed balance, since no active processes are involved:

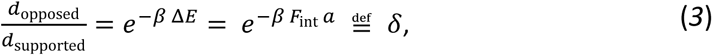

where *β* = 1/*k_B_T*, with *k_B_*-the Boltzmann constant and *T* - temperature. Furthermore, Δ*E* = *F*_int_ *a* is the energy gain associated with moving two interacting particles a distance *a* closer to each other.

The detailed balance constrains only the ratio of the two rates, but not the rates themselves; splitting the Boltzmann factor is thus ambiguous and we have to make a choice. For the sake of simplicity, and motivated by the experimental finding that the velocity of kinesins is typically not or only weakly increased by assisting loads (11, 12), we chose to apply the full Boltzmann factor only to motion away from an interacting particle:

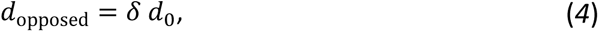

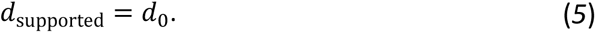

Note that previous studies have suggested that the qualitative behavior of lattice gas models with interacting particles is mostly independent of the specific choice of splitting the Boltzmann factor (13-18).

The functional relation between the rate of active motion and supporting or opposing forces was motivated by the experimentally known velocity-force relation of kinesin-1 (14-18). In agreement with these experimental findings, we assume that active motion is slowed down linearly by forces that oppose this motion, but not or only weakly accelerated by forces that support motion. Specifically, for active minus-end-directed motion of an interacting (denoted by the subscript “int”) particle, we assumed the relations:

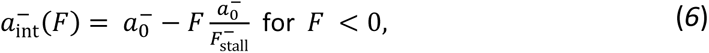

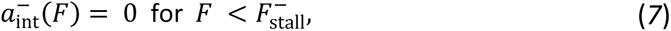

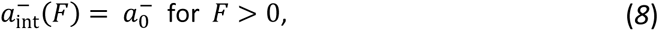

where a negative sign of the force corresponds to a direction opposing motion, i.e., plus-end-directed forces and vice versa, and 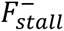 denotes the stall force for active minus-end-directed motion. Note that, in accordance with our convention, the stall force shows a negative value. Analogously, for active plus-end-directed motion, we assume the relations:

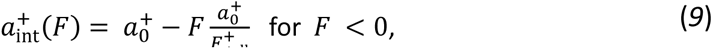

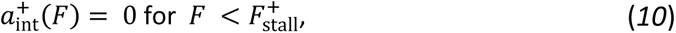

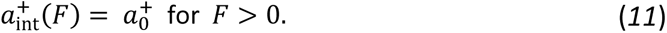

Again, a negative sign denotes forces that oppose the direction of motion, i.e., minus-end-directed forces in this case. Moreover, 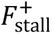 denotes the stall force for active plus-end-directed motion and also shows a negative value. An important assumption required to obtain directionality switching is an asymmetric response of active motion to forces that oppose this motion. Specifically, while minus-end-directed motion is more likely than plus-end-directed motion of a particle for weak or no opposing forces, plus-end-directed motion is more likely for large forces that oppose motion, referred to here as drag. To incorporate such a transition for the preferred directionality of active motion of particles into our model, we assume 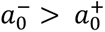 but 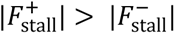. A plot illustrating the resulting velocity-force relation assumed in the model for Cin8 is shown in Fig. SB2 (see next page), which is a reproduction of Fig. 2C, lower panel, of the main text.

Taking Brownian motion and active motion together, we then obtain four different rates for motion of a particle subject to an interaction with another particle:

a. For a plus-end-directed hopping subject to a plus-end-directed (supporting) force:

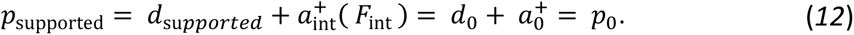
b. For a minus-end-directed hopping subject to a minus-end-directed (supporting) force:

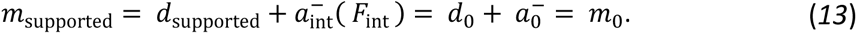
c. For a plus-end-directed hopping subject to a minus-end-directed (opposing) force:

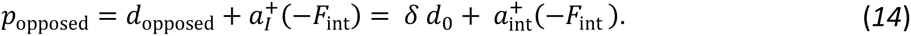
d. For a minus-end-directed hopping subject to a plus-end-directed (opposing) force:

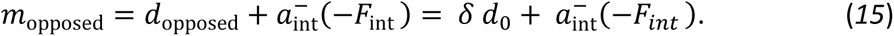

**Figure SB2.**
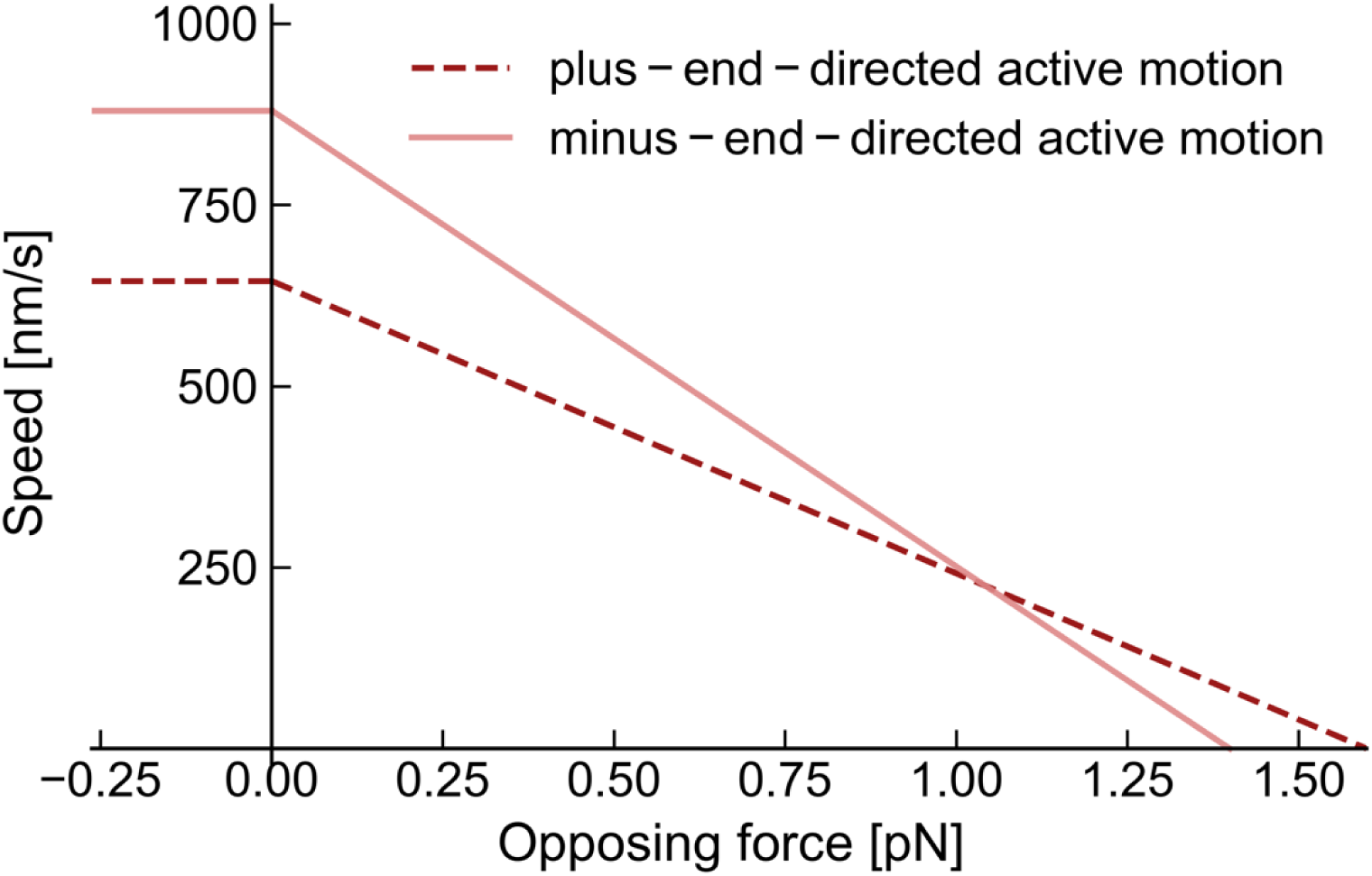
Asymmetric response of active motion of modeled Cin8 motors in the plus-end and minus-end directions to forces that oppose this motion. We assumed that the active motion of particles occurs in both directions on the MT. The responses of active motion in the plus-end and minus-end directions to an external force were asymmetric: Minus-end-directed motion was assumed to be faster at vanishing external forces but to exhibit a smaller stall force (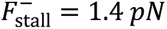) than plus-end-directed active motion (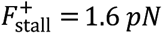). Active motion was not accelerated by a force that acts in the direction of motion (here negative signs) in the model. Note that in this Figure we show the response of active motion to an opposing force, implying that here a positive sign denotes forces acting against the respective direction of motion, whereas a negative sign denotes forces that act in the same direction. The plot is reproduced from Fig. 2C, lower panel, of the main text for convenience.

##### B1.5 Attachment and detachment of attracted particles

We assume that the attachment and detachment of Cin8 is ATP-independent and, hence, constrained by a detailed balance. Consistent with our convention chosen above, we split the Boltzmann factor such that motion away from an interacting particle is weighted by the full Boltzmann factor *e*^−*β* Δ*E*^. Correspondingly, the rate of detachment is reduced for interacting particles, while the rate of attachment is unchanged. In detail, the detachment rate 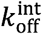 of a particle that interacts with one or two other particles to the left or right, respectively, may be expressed as:

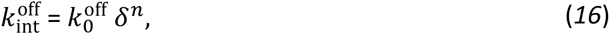

with

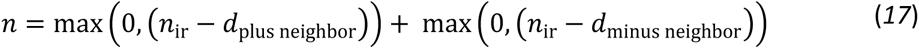

where *n*_ir_ = *R*/*a* is the interaction range in units of the lattice spacing and *d*_plus neighbor_ and *d*_minus neighbor_ are the distances to the neighboring particles in the plus- and minus-end directions, respectively.

An illustration of the model is shown in Fig. SB3.

**Figure SB3.**
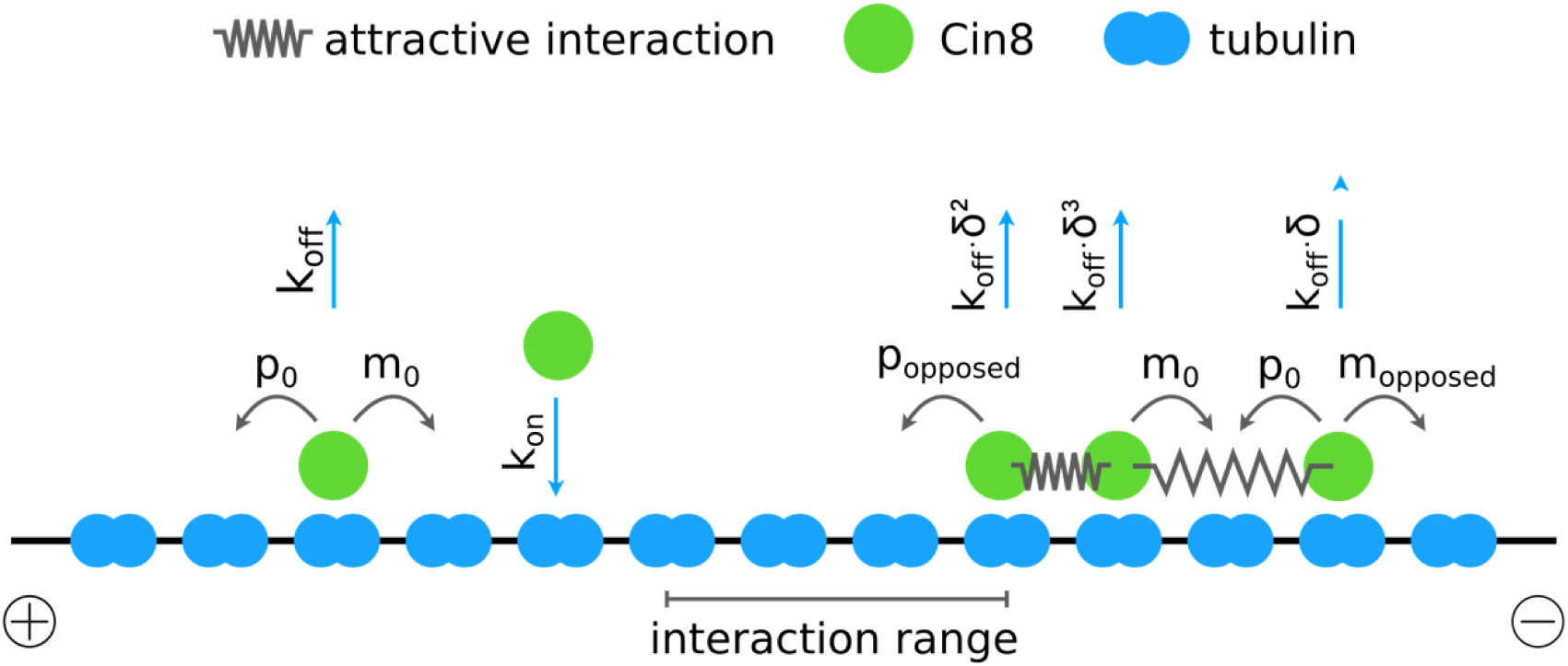
Illustration of the computational model. Particles (green disks) that represent Cin8 motors move stochastically on a discrete, one-dimensional lattice (blue – schematic tubulin dimers) with a lattice spacing of *a* = 8.4 nm that represents the MT. Non-interacting particles move to vacant neighboring lattice sites in the plus-end and minus-end directions at rates *p*_0_ and *m*_0_, respectively, and detach at rate *k*_off_. Particles attach to vacant lattice sites at rate *k*_on_. Particles are subject to attractive interactions (illustrated as springs) with an interaction potential as shown in Figure SB1 for interparticle distances smaller than the interaction range *R* = 3*a*. The motion of particles in the plus-end and minus-end directions that is opposed by the force due to an interaction with another particle occurs at decreased rates *p*_opposed_ and *m*_opposed_, respectively. The motion of particles that is supported by a force is not accelerated. For the sake of simplicity, particles interact only with nearest neighbors. Due to the specific form of interactions as illustrated in Figure SB1 (i.e., a constant interaction force for distances < *R*), a particle that interacts with another particle in the plus-end direction as well as with another particle in the minus-end direction (e.g., second particle from the right in the illustration) effectively moves as a non-interacting particle, since the interaction forces in the two directions cancel out. Interacting particles detach at rate *k*_off_ · *δ^n^* where *n* = max(0, *n_ir_* − *d*_minus neighbor_) + max(0, *n_ir_* − *d*_plus neighbor_). Here, *n_ir_* = *R*/*a*=3 is the interaction range in units of the lattice spacing and *d*_minus neighbor_ and *d*_plus neighbor_ denote the distances to the neighboring particle in the minus-end and plus-end directions (also in units of the lattice spacing), respectively. In this way, detachment fulfills detailed balance and is hence implicitly assumed to occur independently of ATP turnover.

##### B1.6 Simulation algorithm

Simulations of the model were performed using a variant of the Gillespie algorithm (19). The Gillespie algorithm is a stochastic simulation algorithm that produces a (stochastically exact) realization for the temporal evolution of the lattice occupation. We implemented a variant of the direct method, which, in essence, consists of two main steps. In the first step, the waiting time for the next reaction is selected from the exact waiting time distribution. Since we assume Markovian dynamics and time-independent transition rates, the corresponding distribution is an exponential distribution with mean 1⁄*α*, where *α* is the sum of the rates of all particle transitions that are possible in the current configuration. In the next step, one of the possible transitions is selected, and the probability to select a given transition is proportional to the respective rate. To increase computational efficiency, we split the second step into two substeps: we grouped all transition events with identical rates and first selected the type of transition that occurs (i.e., we selected a transition group) and thereafter we selected one of the particles in this group (i.e., we selected a particle in the respective transition group).

##### B1.7 Version of the model that accounts only for the diffusive module of Cin8 motion

As stated in the main text, we used a simplified version of the model that neglects active motion and accounts only for the diffusive module of Cin8 motion at several stages. This simplified version of the model is identical to the one described above, except that all transition rates related to active motion were set to zero, i.e., 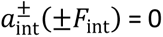 and 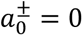.

#### B2 Model parameters

##### B2.1 Determination of the rate corresponding to Brownian motion

The rate *d*_0_, which corresponds to bidirectional ATP-independent motion due to random thermal forces (i.e., Brownian motion) was inferred from the diffusion coefficient and displacement velocity of single (non-interacting) particles measured experimentally. Very generally, the average displacement velocity *v* and diffusion coefficient *D* of a particle that moves stochastically on a lattice at rates *p* and *m* to left- and right-neighboring lattice sites (random walker), respectively, is given by:

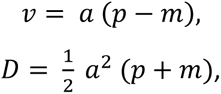

where *a* – the lattice spacing, which, in our case, corresponds to the size of a tubulin dimer of approximately 8.4 nm (10). Inserting in the above equations the diffusion coefficient and displacement velocity of non-interacting Cin8 motors as determined experimentally for single Cin8 molecules (Table 1, main text), we obtain *m* = 1460 s^−1^ and *p* = 1432 s^−1^.

As it is not clear which fraction of these values derives from active motion and which fraction from Brownian motion, we examined typical ATP-turnover rates of other kinesins. At saturating ATP concentrations, kinesin-1 exhibits a maximal velocity of approximately *v*_max_ = 0.9 µm/s (18, 20). Since kinesin-1 hydrolyzes one ATP molecule per step (21, 22), the maximal ATP turnover rate is approximately equal to 110 ATP molecules per second. This rate is one order of magnitude lower than the rates for bidirectional motion computed above. In addition, the diffusion coefficient of 102·10^3^ nm^2^/s as measured for Cin8 is of the same order of magnitude as the diffusion coefficients of other kinesins due to ATP-independent Brownian motion (23-25). Thus, it seems very likely that the largest contribution to the rates of bidirectional motion of Cin8 stems from Brownian motion. Based on this line of argument, and for the sake of simplicity, we assumed that the symmetric component of motion, i.e., a rate of 1432 s^−1^ in both directions, derives solely from Brownian motion, and we correspondingly set *d*_0_ = 1432 s^−1^.

##### B2.2 Determination of the interaction range

**Figure SB4.**
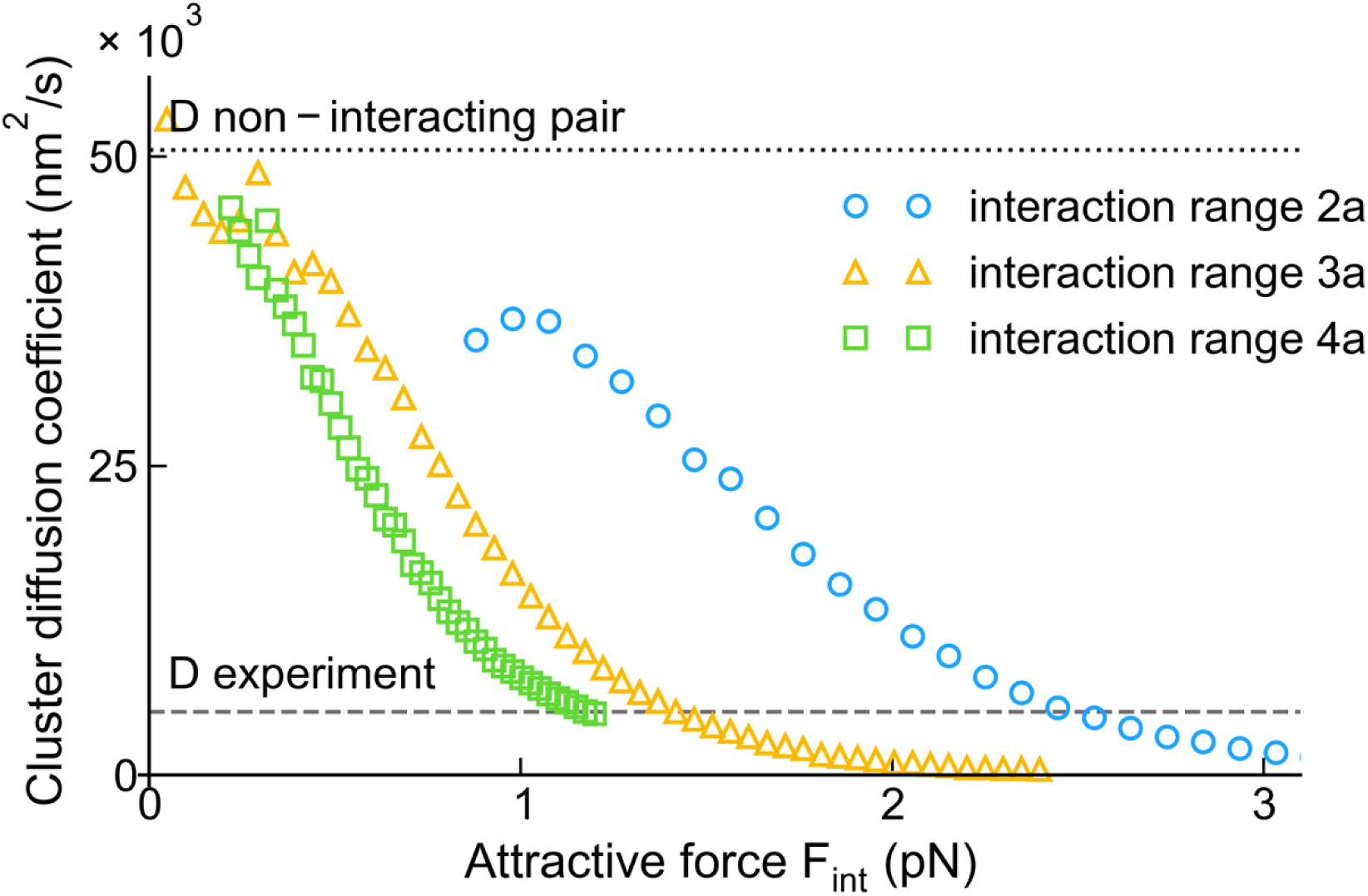
Diffusion coefficient for a pair of particles for different interaction forces between particles and different interaction ranges. The plot shows the diffusion coefficient of pairs of particles obtained from simulations as a function of the interaction force *F*_int_. The analysis allowed us to determine the strength of the attractive forces that led to a diffusion coefficient compatible with that measured in the experiments (dashed gray line): approximately 1.2, 1.4, and 2.5 pN for *R* = 2*a*, *R* = 3*a*, and *R* = 4*a*, respectively. The theoretical value for the diffusion coefficient of a non-interacting pair of motors (dotted line) is shown as a reference. Other parameter values used in the simulation were *p*_0_ = *m*_0_ = 1432 s^−1^; *p*_opposed_ = *m*_opposed_ = *δ p*_0_ with *δ* = exp(−*βF*_int_ *a*) being the Boltzmann factor and *β* = 1/k*T* (with *T* – temperature and k – the Boltzmann constant); and *a* = 8.4 nm denotes the lattice spacing. A temperature corresponding to *β*^−1^= 4.11 pN nm was used. Attachment and detachment rates were *k*_on_ = 2 · 10^−5^*s*^−1^, *k*_off_ = 0.0625 *s*^−1^, respectively. A lattice size of L = 5000 was used.

To choose an appropriate value for the interaction range *R* in the model, we performed simulations using an interaction range of two, three, or four lattice sites (i.e., 2*a*, 3*a*, and 4*a*). First, in analogy to the main text, we determined the strength of interactions required to reproduce the experimentally measured diffusion coefficient of pairs of particles for each of the values of *R*. As argued in the main text, this quantification can be made by using the model that incorporates only the diffusive module of motion, as the diffusion coefficient of particles is likely to be dominated by Brownian motion. For each of the three different interaction ranges, we determined the relation between the diffusion coefficient for pairs of particles and the interaction strength as shown in Fig. SB4. For interaction ranges of {2*a*; 3*a*; 4*a*} an interaction strength of approximately {1.2; 1.4; 2.5} was found to reproduce the experimentally determined diffusion coefficient of pairs of Cin8 particles (5000 nm^2^/s).

In the next step, we used this quantification of the attractive interactions between particles to perform simulations at particle attachment rates that corresponded to those measured in the experiments. Representative kymographs for *R* = {2*a*; 3*a*; 4*a*} are shown in Fig. SB5(a). While an interaction range of *R* = 2*a* led to the formation of hardly any clusters, an interaction range of *R* = 4*a* led to systems that were dominated by large clusters. In contrast, an interaction range of *R* = 3*a* caused the formation of clusters that were stable but yet did not dominate the lattice. This qualitative observation was further supported by the distribution of the number of particles in a cluster measured in simulations for the different interaction ranges as shown in Fig. SB5(b). Unlike the behavior for interaction ranges of size *R* = {2*a*; 4*a*}, an interaction range of *R* = 3*a* produced clusters that exhibited sizes and lifetimes compatible with experiments. Taken together, this stability analysis of clusters suggests that only an interaction range of *R* = 3*a* agrees with experimental data, while interaction ranges of *R* = {2*a*; 4*a*} produced unstable systems in which clusters either hardly formed at all or dominated the system.

**Figure SB5.**
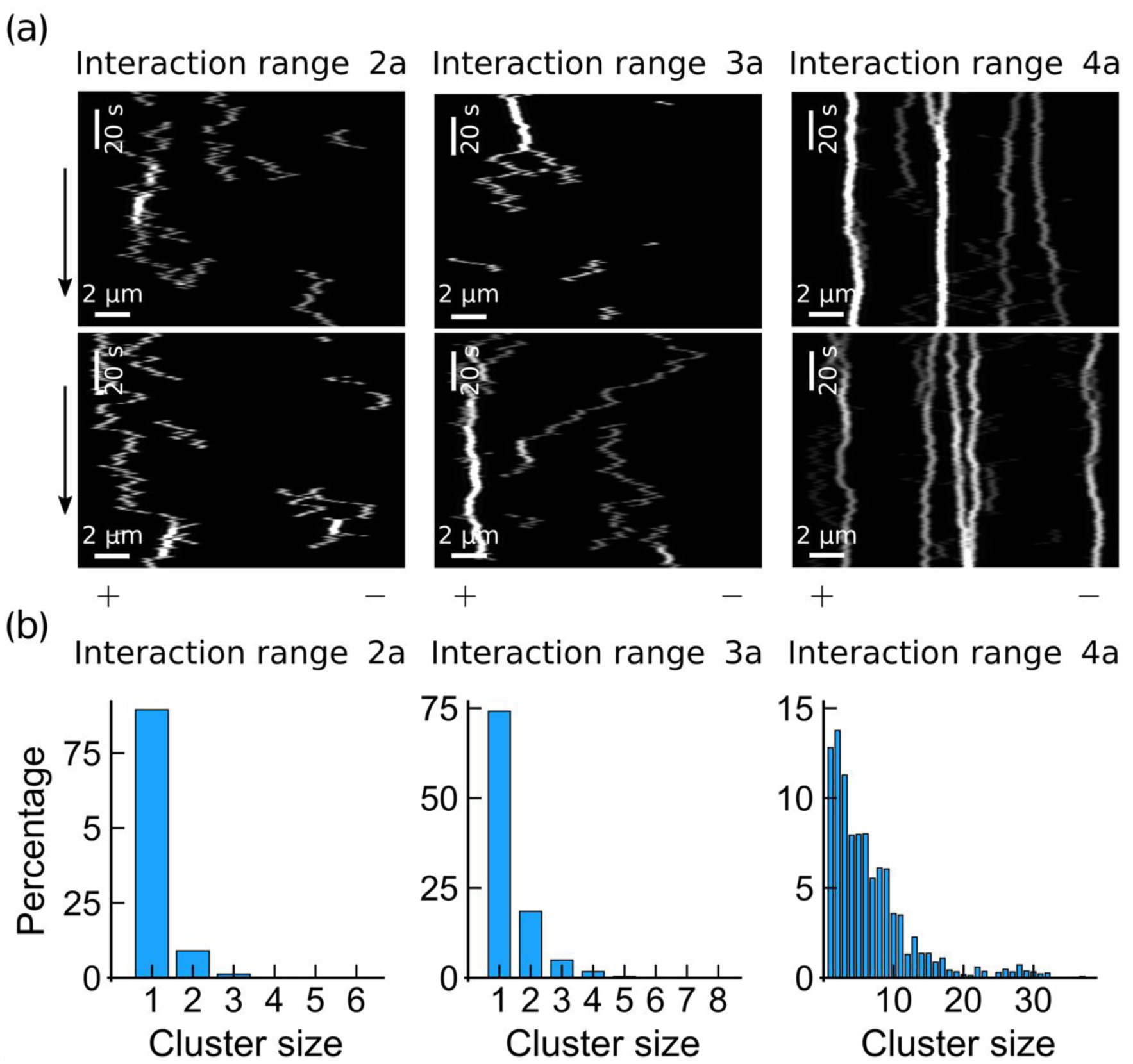
Analysis of the number of particles in a cluster (cluster size) for different interaction ranges *R*. **(a)** Illustrative kymographs obtained from simulations of the model that incorporates only the diffusive module of particle motion for *R* = {2*a*, 3*a*, 4*a*}. Only for interaction ranges of *R* = 3*a* was the formation of clusters significant yet not dominating; for *R* = 2*a* clusters hardly formed at all, and for *R* = 4*a* the formation of large clusters dominated the system. **(b)** The probability distributions of the cluster size (in %) as obtained from simulations with *R* = {2*a*, 3*a*, 4*a*} support the notion that the formation of stable but small clusters is likely only for *R* = 3*a*. Other parameters in **(a)** and **(b)** were *p*_0_ = *m*_0_ = 1432 s^−1^, *p*_opposed_ = *m*_opposed_ = {9.6; 79; 130} *s*^−1^ for *R* = {2*a*, 3*a*, 4*a*}, respectively, *k*_on_ = 4.8 · 10^−5^ *s*^−1^, and *k*_off_ = 0.0625 *s*^−1^. The corresponding Boltzmann factors consistent with the rates of motion against an opposing force that were used to compute the reduced detachment rates of interacting particles were *δ* = {6.7 · 10^−3^; 5.5 · 10^−2^; 9 · 10^−2^} *s*^−1^(12) The lattice size was set to L = 2000 in **(a)** and **(b)**. Plus and minus signs denote the respective lattice ends. Black arrows indicate the direction of time. The percentage of the respective cluster size in **(b)** was computed by relating the summed existence times of all single particles and clusters for each of the different size categories.

##### B2.3 Determination of the rate corresponding to active motion

To specify active motion of particles in our model, we first defined a force-velocity relation for active plus- and minus-end-directed motion that is compatible with experimental observations. Specifically, the relation is constrained by the approximate values for the stalling forces measured in previous experiments, on the one hand, and by the average displacement velocity of non-interacting Cin8 motors measured experimentally in our study, on the other. In the following text, we discuss both of these constraints.

The stall force for Cin8 was recently inferred from pushing forces of Cin8 on gliding MTs in a motility assay (26); for both directions, a stall force of approximately 1.5 pN was found. Based on the argument provided in the Model Description, we assumed that 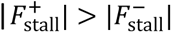. To comply with these constraints, we chose 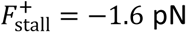 and 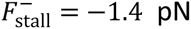. This implies that 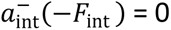.

The average displacement velocity of non-interacting Cin8 motors of *v* ≈ −237 nm/s (see Table 1 of the main text) further constrains the force-velocity relation of plus-end and minus-end-directed motion of Cin8. Specifically, without opposing forces, the difference between active plus-end-directed motion and minus-end-directed motion must equal the measured displacement velocity, i.e., 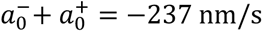.

**Figure SB6.**
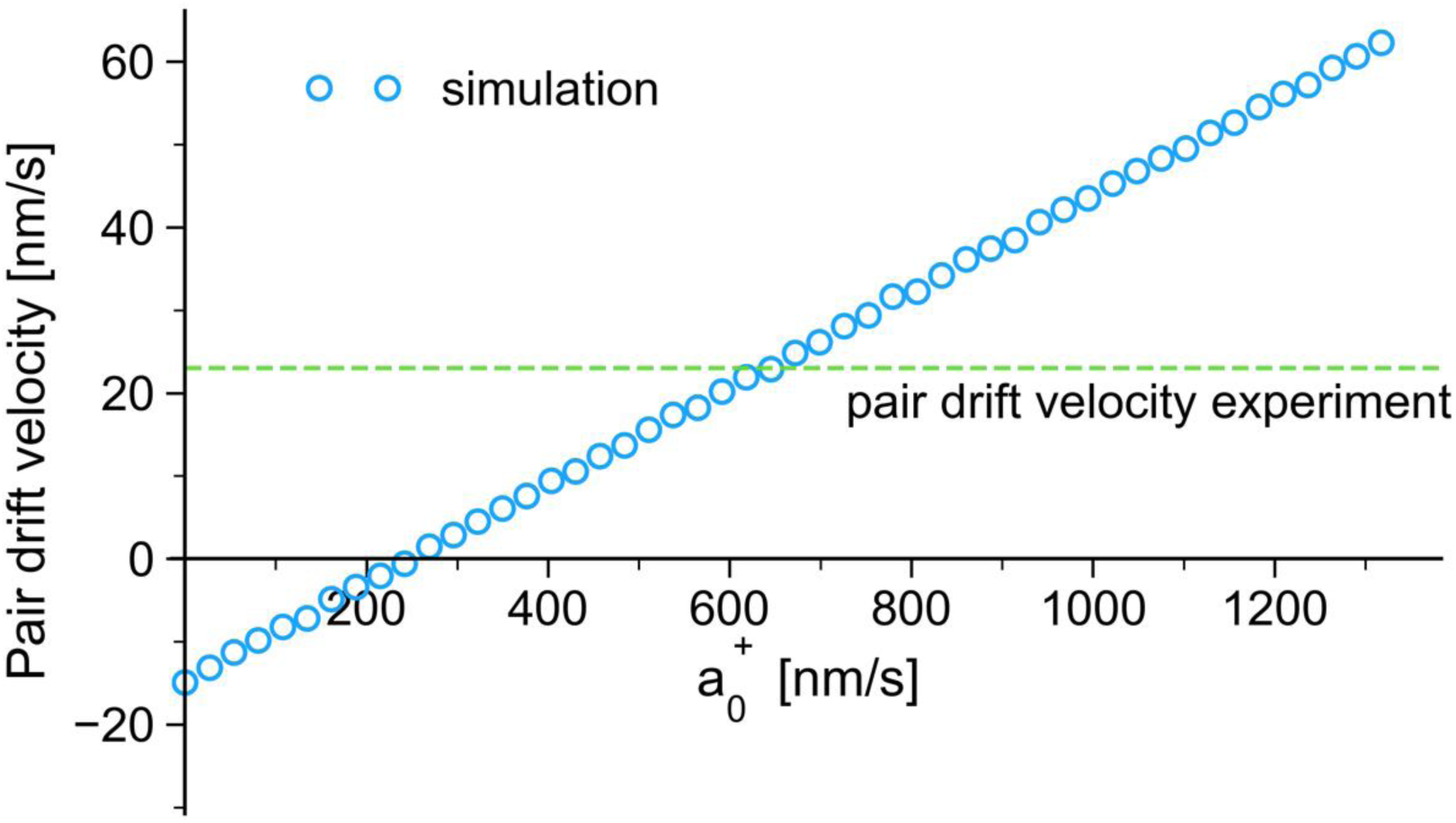
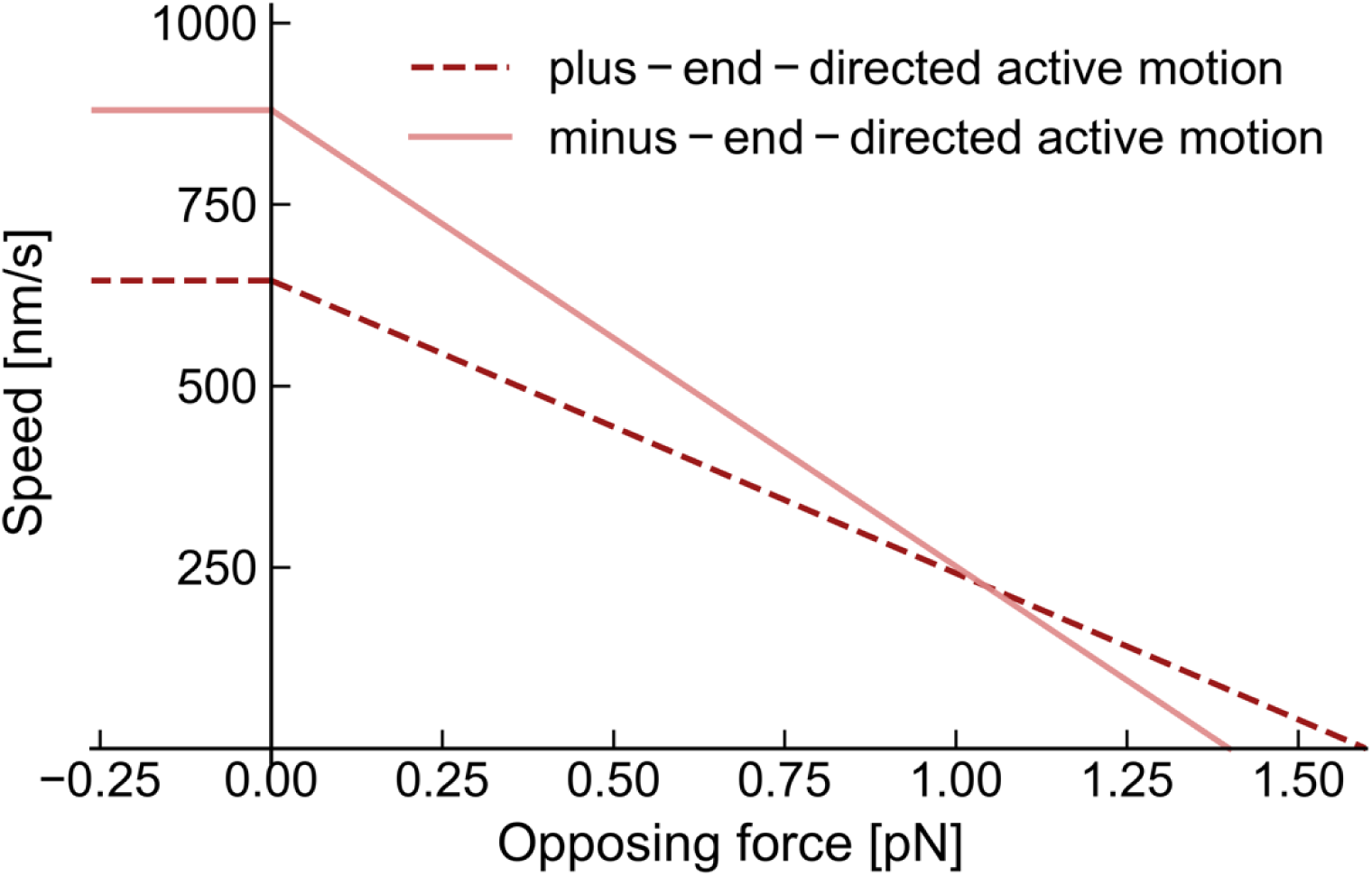
Relationship between the drift velocity of a particle pair and the rate 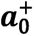. Simulation results indicate a linear relation between the rate 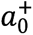 of an individual particle and the drift velocity of a pair of particles. A drift velocity of a pair of Cin8 motors as determined in experiments (dashed line) is reproduced by a value of 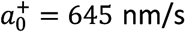 in the model. Particle dynamics were chosen to fulfill the linear force-velocity relation and stall forces (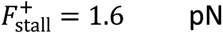, 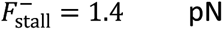) as specified in Figure SB2, as well as a drift velocity of approximately −235 nm/s for a non-interacting particle. Note that we assumed that the force related to particle interactions stalls active motion in the minus-end direction, 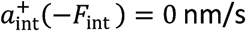. The other parameter values used in the simulation were *d_0_* = 1432 s^−1^, *d*_opposed_ = 78.8 s^−1^, 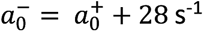, 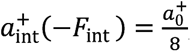, 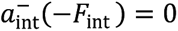, *k*_on_ = 2 · 10^−5^s^−1^, *k*_off_ = 0.0625 s^−1^. Simulated lattice size was *L* = 5000. Here, negative signs of velocities indicate minus-end-directed motion. Moreover, negative signs of forces denote forces that oppose motion while positive signs denote forces that support motion.

Based on the above constraints, we then systematically varied the velocity 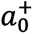 of active minus-end-directed motion in the absence of an external force in the simulations, and determined the average displacement velocity of pairs of particles. The resulting relation between the average displacement velocity and 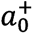 is shown in Fig. SB6 and yields a value of approximately 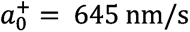 and, consequently, 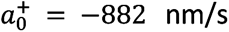. With the constraints stated above, these values correlate with speeds of approximately 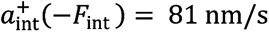 and 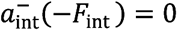 when active plus- and minus-end-directed motion is opposed by the force of a particle interaction, respectively.

##### B2.4 Determination of attachment and detachment rates

The detachment rate was inferred from the average dwell time of 16 ± 2 s of individual Cin8 motors measured experimentally. The attachment rate of Cin8 particles was also measured at 1–2 pM, and yielded a value of 5.74 ± 0.68 · 10^−3^ μm^−1^ s^−1^, which was converted to the attachment rate per lattice site of size *a* = 8.4 nm: 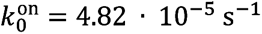. Together with the strength of an attractive force of 1.4 pN between interacting Cin8 motors, as determined in the main text, the above values yielded a completely quantified model without any free parameters. The resulting list of dynamic model parameters is given in Table SB1. Table SB2 provides an overview of the mechanistic interpretations of the model parameters.

**Table SB1.** Summary of the parameters for the computational model.

| Parameter | Value |
| --- | --- |
| Interaction range $R$ | 3 lattice sites |
| Minus-end-directed motion; non-interacting particles $m_0$ | $1.5368 \cdot 10^3 \text{ s}^{-1}$ |
| Plus-end-directed motion; non-interacting particles $p_0$ | $1.5088 \cdot 10^3 \text{ s}^{-1}$ |
| Minus-end-directed motion against the force of interactions $m_{opposed}$ | $7.88 \cdot 10^3 \text{ s}^{-1}$ |
| Plus-end-directed motion against the force of interactions $p_{opposed}$ | $8.84 \cdot 10^3 \text{ s}^{-1}$ |
| Attachment $k_0^{on}$ | $4.82 \cdot 10^{-5} \text{ s}^{-1}$ |
| Detachment; non-interacting particle $k_0^{off}$ | $6.25 \cdot 10^{-2} \text{ s}^{-1}$ |
| Boltzmann factor $\delta$ | $5.5 \cdot 10^{-2} \text{ s}^{-1}$ |

The Boltzmann factor is used in simulations to weight the detachment rate of interacting particles. Specifically, the detachment rate is multiplied by $\delta^n$ , where $n = \max(0, (n_{ir} - d_{plus \text{ neighbor}})) + \max(0, (n_{ir} - d_{minus \text{ neighbor}}))$ . Here, $n_{ir}$ denotes the interaction range in units of lattice sites and $d_{plus \text{ neighbor}}$ and $d_{minus \text{ neighbor}}$ denote the distances to the neighboring particle in the plus-end and minus-end directions in units of lattice sites, respectively.

**Table SB2.** Overview of several model parameters in terms of physical quantities.

| Parameter | Value |
| --- | --- |
| Interaction range | 25 nm |
| Interaction strength | 5.8 $k_B T$ |
| Force between interacting particles | 1.4 pN |
| Speed of plus-end directed active motion when opposed by a particle interaction | 81 nm/s |
| Speed of minus-end directed active motion when opposed by a particle interaction | 0 nm/s |

A temperature corresponding to $k_B T = 4.11 \text{ pN nm}$ was implicitly assumed.

#### B3 Cluster tracking in simulations

In simulations, a cluster was defined as a group of particles with mutual interparticle distances below the resolution limit of approximately 400 nm. This translates to a maximal distance 48 lattice sites between two neighboring particles in a cluster. Note that this distance substantially exceeds the interaction range of three lattice sites.

In analogy to experiments, we chose a temporal resolution of 1.25 s for cluster tracking in simulations, i.e., we evaluated the positions of single particles and clusters at intervals of 1.25 s.

#### B4 Mean displacement and mean squared displacement analyses of the quantified model

To test whether the fully quantified model produces displacement velocities and diffusion coefficients of single particles and clusters that agree with those measured *in vitro* for Cin8 motors, we performed MD and MSD analyses of particle and cluster trajectories obtained from simulations of the computational model. Table SB3 presents a list of the ensuing average displacement velocities obtained from the MD (*v*_*MD*_) and MSD analyses (*v*_*MSD*_), as well as the diffusion coefficients (*D*_*MSD*_) obtained from the MSD analysis. All quantities showed excellent agreement with the experimental data, as shown in Table SB4 (which is reproduced from Table 1 of the main text for convenience).

**Table SB3.** List of the drift velocities and diffusion coefficients of single Cin8 particles and clusters in the fully quantified model for the collective motion of Cin8.

| Simulation | $v_{MD}$ [nm/s] | $v_{MSD}$ [nm/s] | $D_{MSD}$ [$10^3$ nm/s] |
| --- | --- | --- | --- |
| Single particles | -229 | 227 | 109 |
| Pairs of particles | 23.0 | 22.0 | 5.30 |
| Clusters with three particles | 17.8 | 16.5 | 2.38 |
| Clusters with four particles | 11.5 | 9.58 | 1.79 |
| Clusters with five particles | 9.86 | 8.25 | 1.32 |

Values were obtained by fitting the linear function $\langle x(t) \rangle = v_{MD}t$ to the temporal evolution of the mean displacement, and the quadratic function $\langle x(t)^2 \rangle = v_{MSD}^2 t^2 + 2 D_{MSD}t$ , to the temporal evolution of the mean squared displacement, where $x(t)$ denotes the position of the respective particles at time $t$ on the MT and angle brackets refer to an ensemble average over different trajectories. An attachment rate of $k_0^{on} = 2 \cdot 10^{-5} \text{ s}^{-1}$ was used. Other model parameters are listed in Table SB1. A lattice size of $L = 5000$ was used.

**Table SB4.**
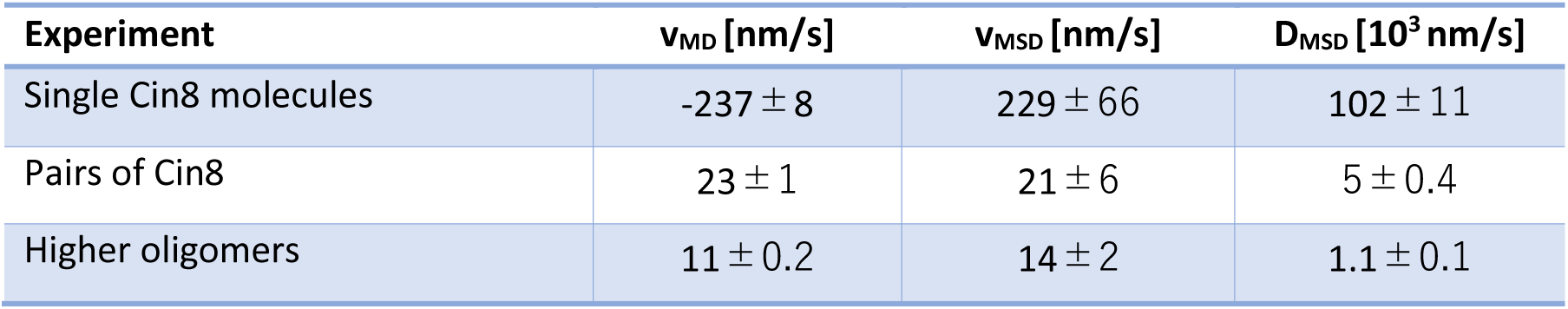
List of the displacement velocities and diffusion coefficients of single Cin8 molecules, pairs of Cin8, and Higher oligomers with more than two motors as obtained by a MD and MSD analyses of the experimental data. The table is a reproduction of Table 1 in the main text.

| Experiment | $v_{MD}$ [nm/s] | $v_{MSD}$ [nm/s] | $D_{MSD}$ [$10^3$ nm/s] |
| --- | --- | --- | --- |
| Single Cin8 molecules | $-237 \pm 8$ | $229 \pm 66$ | $102 \pm 11$ |
| Pairs of Cin8 | $23 \pm 1$ | $21 \pm 6$ | $5 \pm 0.4$ |
| Higher oligomers | $11 \pm 0.2$ | $14 \pm 2$ | $1.1 \pm 0.1$ |

#### B5 Concentration-dependent clustering of the model without active motion

As stated in the main text, we used a version of the computational model that neglects active motion of Cin8 and includes only the diffusive module of Cin8 motion to infer the strength of interactions between Cin8 motors from the diffusion coefficient of pairs of Cin8. Neglecting active motion for this purpose is justified since, as argued above, the diffusion coefficient is very likely to be determined by Brownian motion. To show that even this preliminary version of the model can indeed capture the concentration-dependent clustering behavior of Cin8, we performed simulations at three different concentrations and compared the resulting distribution of cluster sizes with the experimental data. The result is presented in Fig. SB7 and shows good agreement between model and experiment.

**Figure SB7.**
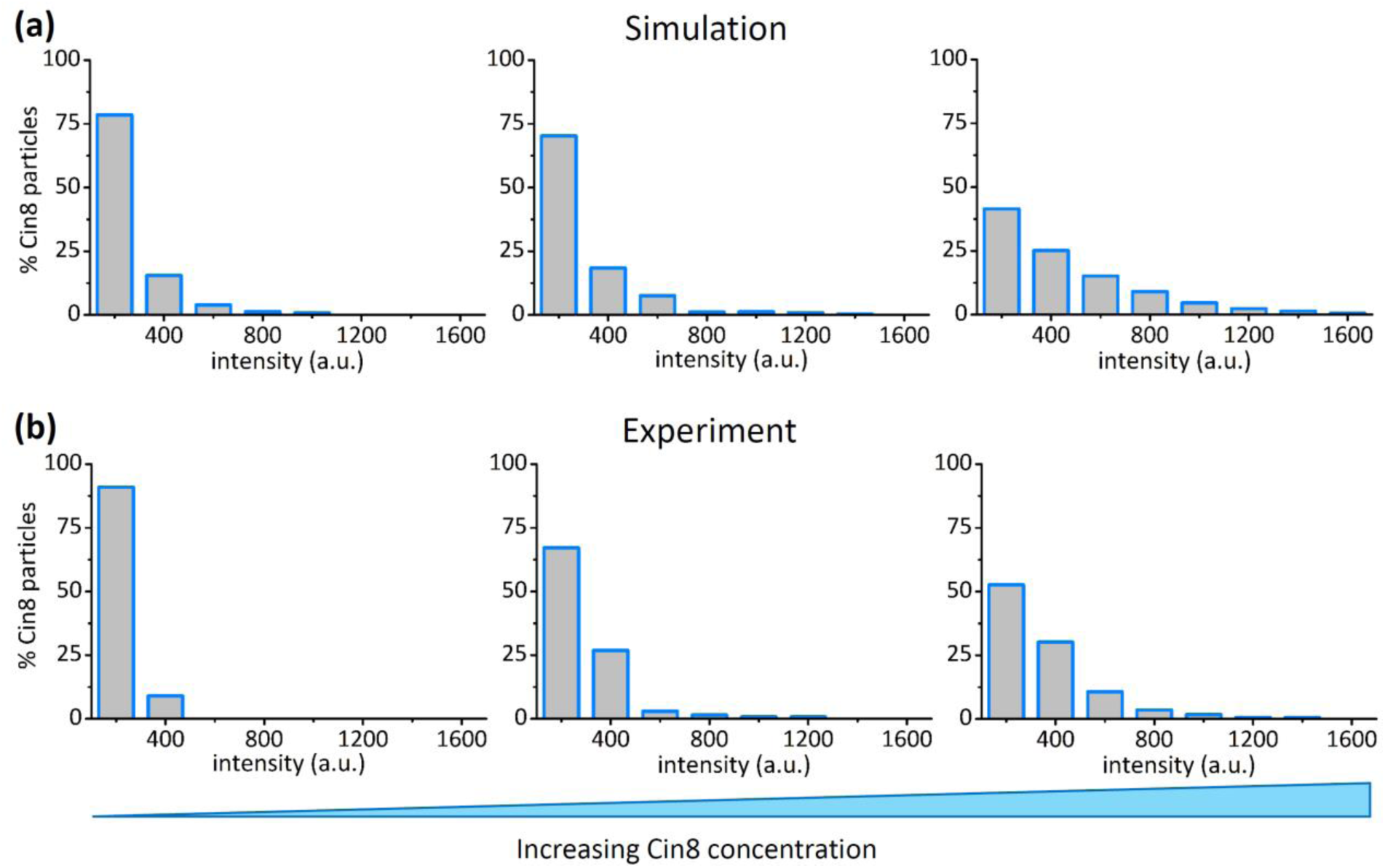
Comparison of the distribution of cluster sizes determined in experiments in the presence of ADP with those obtained in simulations of the model without active motion at different concentrations of Cin8. Both simulations **(a)** and experiments **(b)** consistently showed that for increasing Cin8 concentrations (left to right) very large clusters of Cin8 form and dominate the lattice for high concentrations. In **(a)**, the attachment rates were computed based on the measured attachment rate at medium concentrations (1-2 pM): *k*_on_ = {3.62 · 10^−5^, 4.82 · 10^−5^, 9.04 · 10^−5^} from the left to the right. Lattice size in the simulations was L = 2000; other parameters were as listed in Table SB1. The percentage of the respective cluster size observed in the simulations was computed by relating the summed existence times of all single particles and clusters for each of the different size categories. In **(a)** intensity was determined by multiplying the number of single Cin8 molecules in a cluster by 200 a.u., the maximal intensity of a single Cin8 tetramer. In **(b)**, concentrations were *c* = {0.75 − 1.5, 1 − 2, 1.875 − 3.75} pM from the left to the right.

#### B6 Detailed computation of the caterpillar motion of a pair of Cin8

Below we provide a detailed computation of the motion of a pair of particles to show that plus-end-directed motion of this cluster is more likely than minus-end-directed motion.

**Figure SB8.**
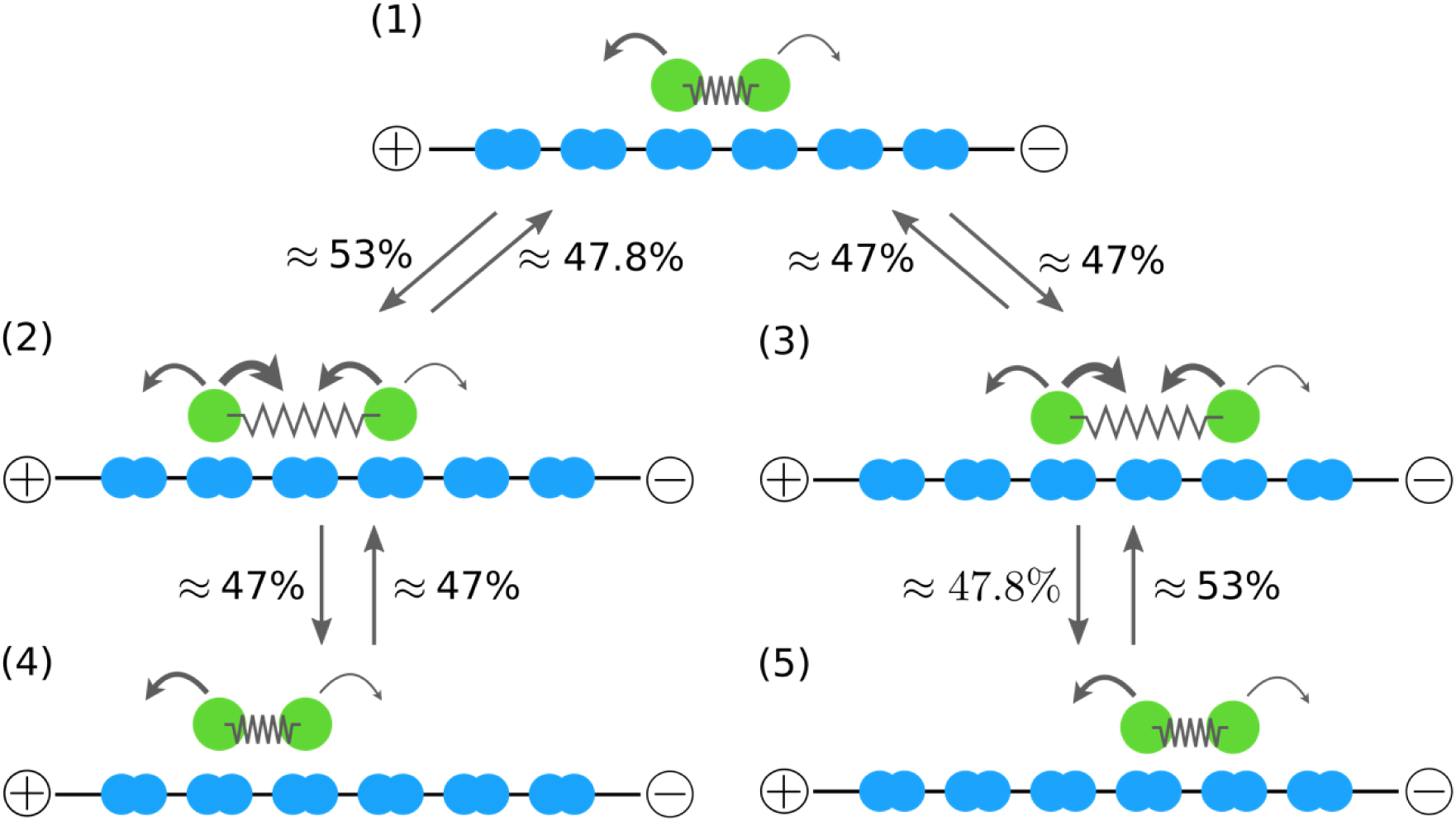
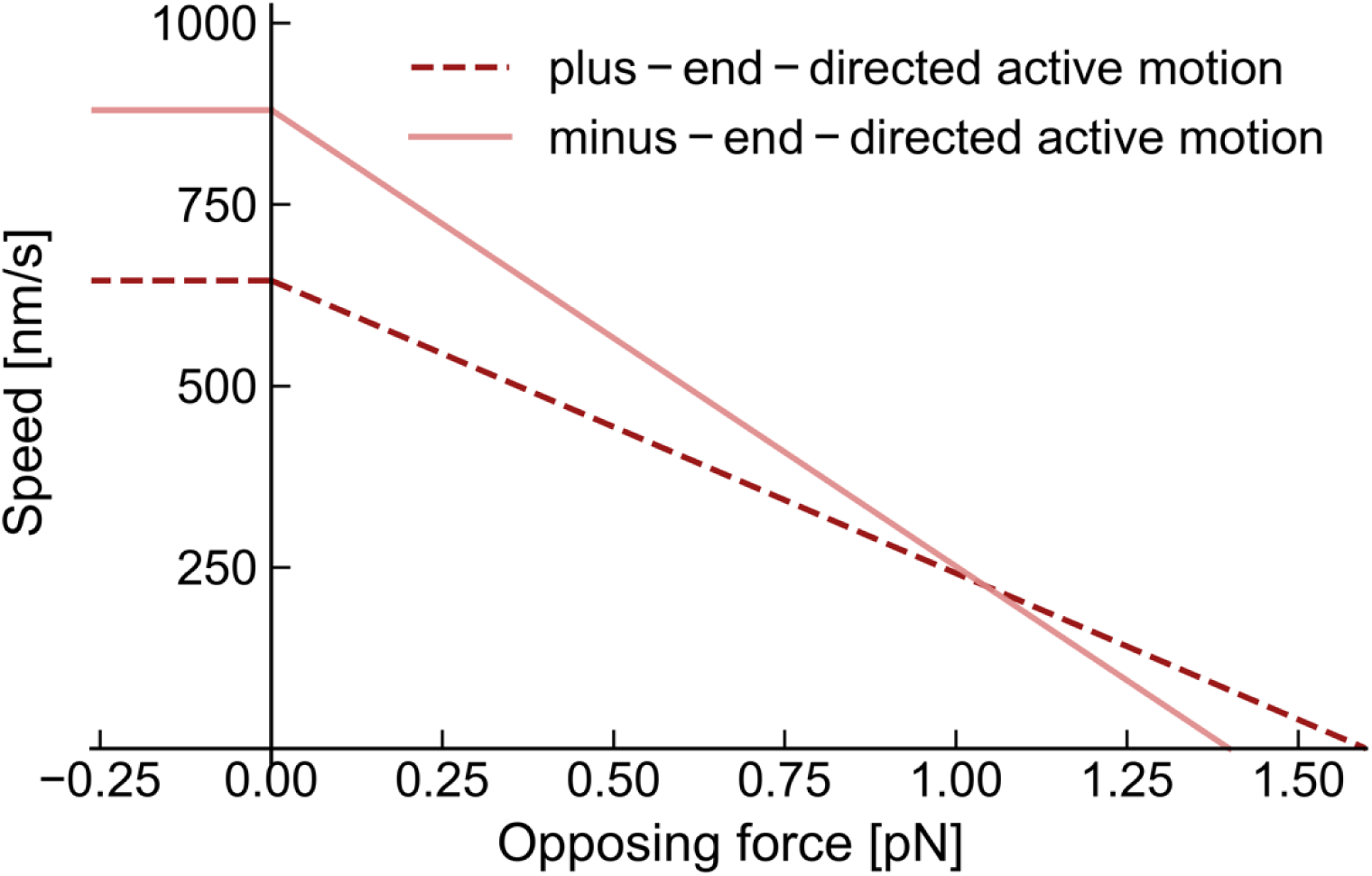
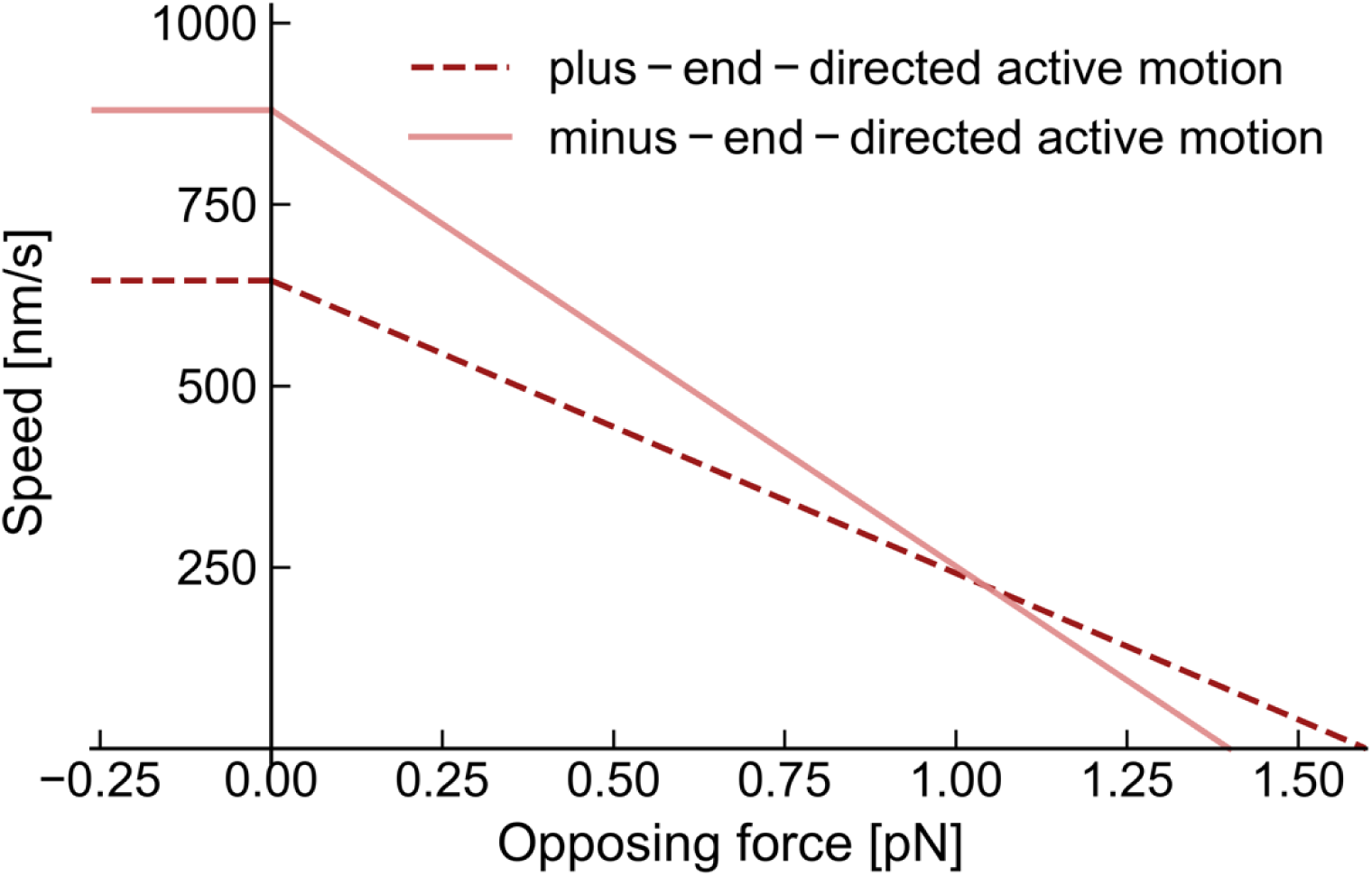
Illustration showing that an asymmetric dependence of active motion on drag biases the motion of particle pairs towards the plus end. Due to the asymmetric response of active motion to drag (see also **Figure SB2**), particles move against an opposing force with a preference towards the plus end. Therefore, a particle pair in a completely compressed state (1) will expand with a slightly higher probability towards the plus end [transition (1) → (2) with probability (53%)] than it does towards the minus end [transition (1) → (3) with probability 47%]. In a cluster configuration where the particles are separated at least by one lattice site, but still interact with each other [e.g., states (2) and (3)], the most likely reaction is motion of the motors closer together (cluster contraction; corresponding probability approximately 95%), which occurs preferentially towards the minus end, since the attractive interaction now supports (rather than opposes) such motion (see also **Figure SB2**). Supporting forces, however, also imply a larger diffusive contribution to the rate of compression of a cluster as compared to the rate of stretching of a cluster (Arrhenius law). As this diffusive component contributes equally to motion of the left and right motors, the probabilities of compression owing to movement towards the minus and the plus ends, respectively, differ only insignificantly (47.8% vs. 47%). As a result, a caterpillar-like motion of a particle pair towards the plus end [transition (1) → (2) → (4)] shows a higher transition probability than motion of a particle pair towards the minus end [transition (1) → (3) → (5)].

Consider two particles on neighboring lattice sites as shown in Fig. SB8(1). The particles may move in one of two different ways: either by minus-end-directed motion of the right particle or by plus-end-directed motion of the left particle. Due to the interaction between the particles, these processes occur at rates *p*_opposed_ for plus-end-directed motion and *m*_opposed_ for minus-end-directed motion. Accordingly, the probability for plus-end-directed motion (conditional on cluster stretching) may be expressed as 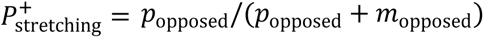. Inserting the values listed in Table SB1 into this equation yields a probability of 53% for the left particle to move towards the plus end, given that cluster stretching occurs. Once the particles will have moved one lattice site further apart by either of the two possibilities, the cluster [now in state (2) or (3)] may either contract (i.e., the particles may move closer together) or expand (i.e., the particles move even further apart). Due to the interaction of the particles, the probability *P*_contraction_ of cluster contraction is substantially larger than that of cluster stretching: *P*_contraction_ = (*m*_supported_ + ^*p*^supported^)/(*m*^supported ^+ *p*^supported ^+ *m*^opposed ^+ *p*^opposed^) ≈ 95%. Thus, for the sake of^ simplicity, we neglect the possibility of further stretching and consider the case in which the cluster contracts again. As active motion of the motors for cluster contraction is supported (and not counteracted) by the motor interaction, the directional biases are the same as those in the absence of external forces, due to our modeling assumptions (see also the section “Description of the stochastic model for Cin8 dynamics”). Hence, hopping of the left particle towards the minus end is favored over plus-end directed motion of the right particle. However, the probability that minus-end-directed motion of the right particle will occur (assuming cluster contraction) is only slightly larger than the probability of plus-end-directed motion of the left particle: 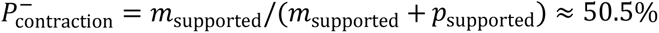. This small difference in probabilities may be attributed to Brownian motion. This form of motility is an identical but mirror-symmetric component of motion of the left and right particles which, in fact, contributes the largest component of motor motion involved in cluster contraction (rate of Brownian motion *d*_0_ = 1432 s^−1^; rates of active motion: 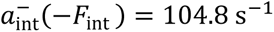 and 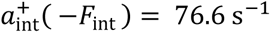 for minus- and plus-end-directed active motion, respectively). Unlike cluster contraction, cluster stretching is opposed by particle interaction, which suppresses diffusion (see also Eq. 3), such that the directional bias is more relevant for cluster stretching than for cluster contraction. As the directional bias of cluster stretching towards the plus end is more pronounced than the directional bias of cluster contraction towards the minus end, it is more likely that the pair of particles will move sequentially towards the plus end [sequence of states (1)->(2)->(4)] than to the minus end [sequence of states (1)->(3)->(5)].

#### B7 Simulation results for reduced Brownian motion

To assess the role of Brownian motion in the directionality switching of Cin8 due to clustering, we performed additional simulations with strongly reduced Brownian motion. Specifically, we lowered the diffusion coefficient of Cin8 motors due to Brownian motion to a value of 128 nm^2^/s in simulations. Representative kymographs of these simulations are shown in Fig. SB9. Motors with such a low diffusion coefficient do not exhibit clustering-induced directionality switching.

**Figure SB9.**
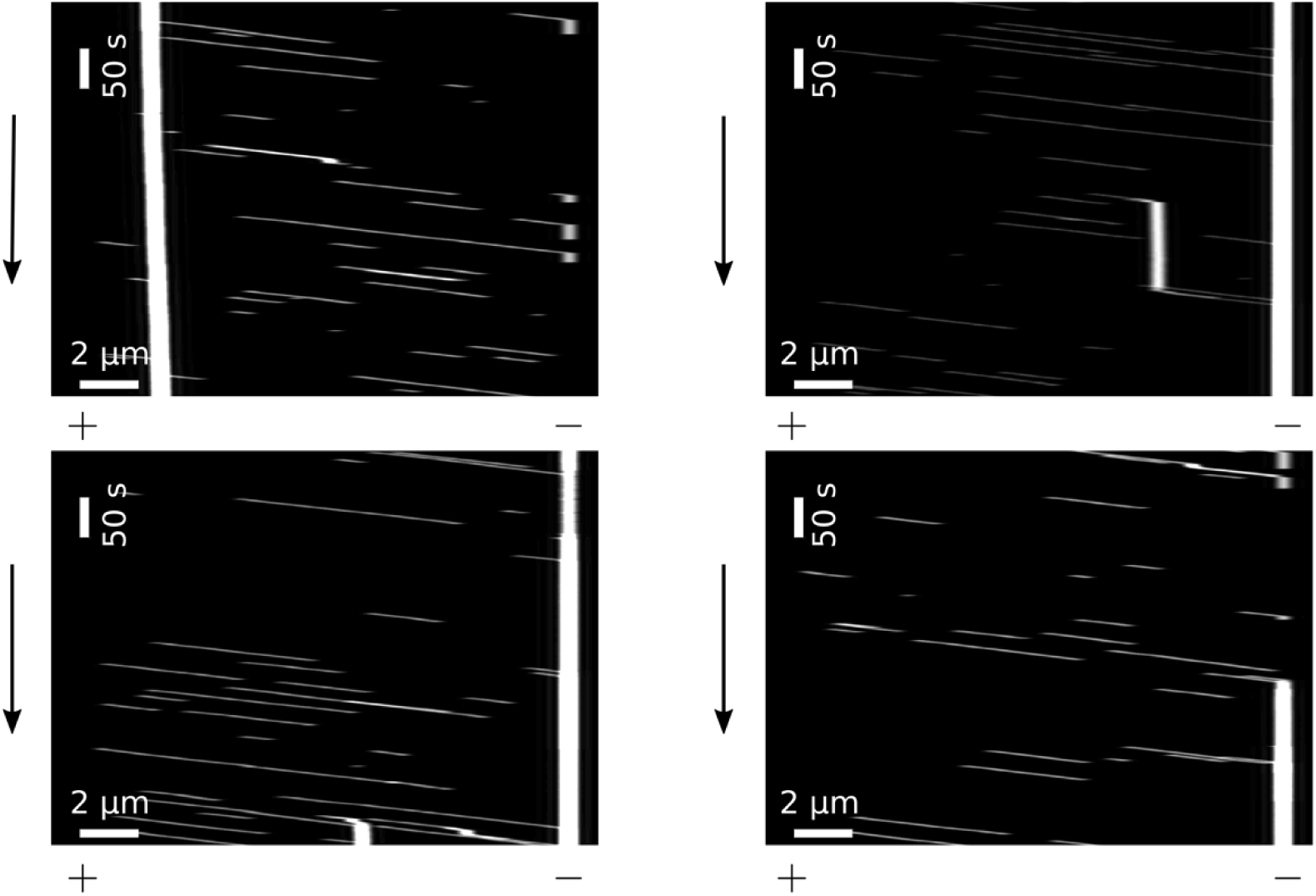
Representative kymographs of simulations with a negligible diffusive component of Cin8 motion. When the diffusive component in the motion of particles was lowered in the model, clusters showed either no net displacement or a slight tendency to move towards the minus end. Plus and minus signs denote the respective lattice end. Black arrows denote the direction of time. Parameter values were *m*_0_ = 29.8 s^−1^, *p*_0_ =1.8 s^−1^, *m*_opposed_ = 0.1 s^−1^, *p*_opposed_ = 9.7 s^−1^, *k*_on_ = 4 · 10^−5^ s^−1^, *k*_off_ = 0.0625 s^−1^. The lattice size was *L* = 2000.

#### B8 Onset of clustering in simulations

When increasing the concentration of Cin8 (i.e., increasing the Cin8 landing rate) in simulations, we noted a very rapid transition from the absence of clusters at low concentrations to a very pronounced clustering at high concentrations. Above a certain Cin8 concentration, clusters were so stable that they continuously accumulated more and more particles, leading to a growing occupational density of particles on the lattice (representing the MT). Fig. SB10 shows the average occupation of the lattice by particles averaged over the whole lattice. As the densities on the lattice increased only very slowly, we chose very long simulation times that exceeded those feasible in experiments by several orders of magnitude. The rapid onset of cluster formation in simulations occurred at attachment rates of approximately 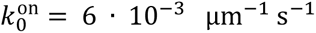. Strikingly, this attachment rate coincides very well with that measured in the experiments, namely, approximately 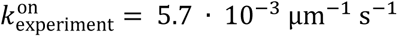 at concentrations at which the formation of clusters was empirically found to begin.

**Figure SB10.**
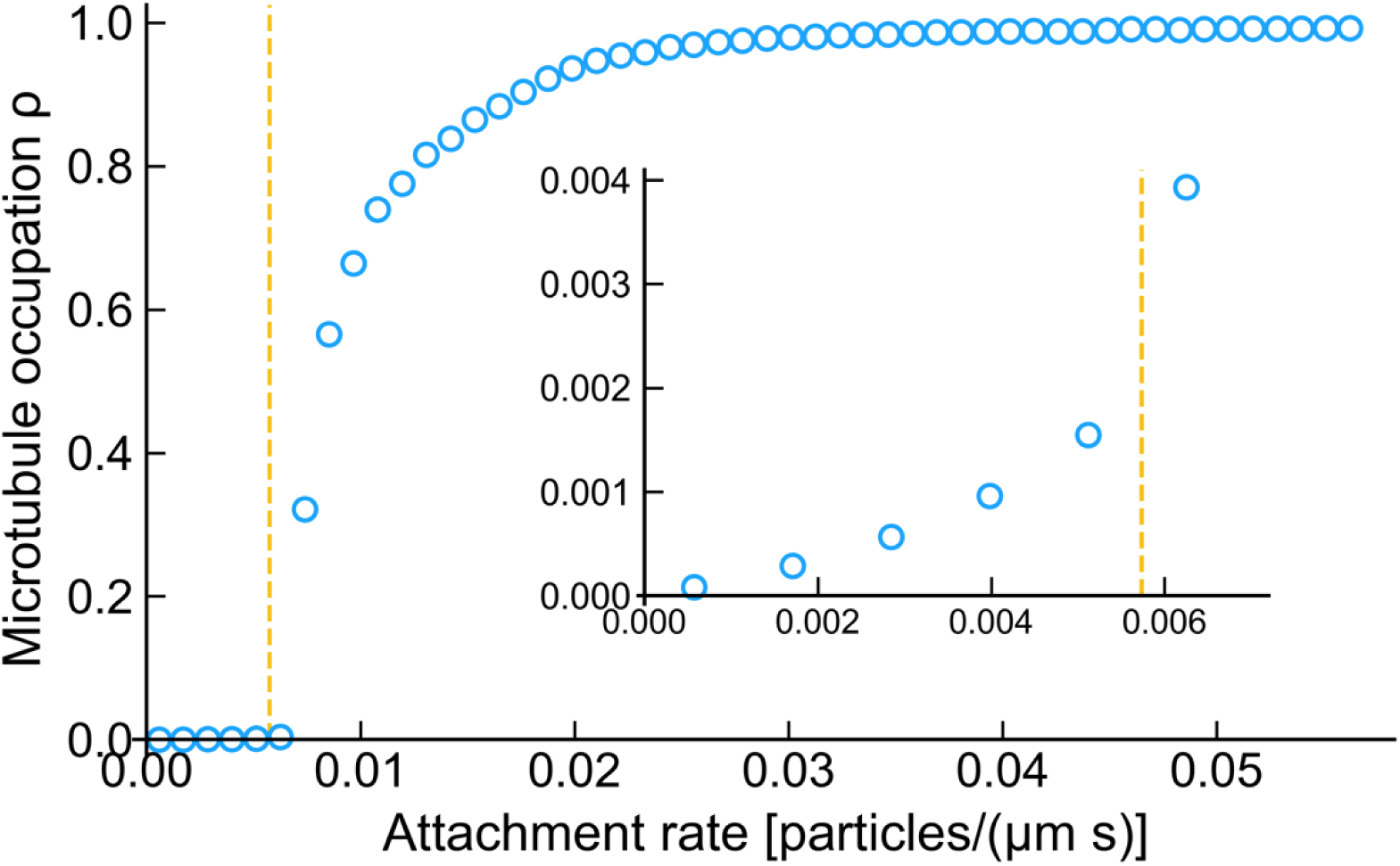
Average occupation of the lattice obtained from simulations with varying particle attachment rates. When the attachment rate of particles was increased—which corresponds to increasing particle concentrations in the experiment—the average occupation of the lattice by particles *ρ* showed an abrupt change at attachment rates of approximately *k*_on_ = 6 · 10^−3^ µm^−1^s^−1^. For larger attachment rates, clusters tended to grow steadily until almost the whole lattice was occupied. Cluster growth in simulations became significant for attachment rates very close to those measured in the experimental setup (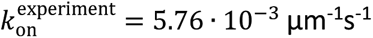, dashed line) at concentrations of c = 1–2 pM. At these concentrations, clustering also became significant in the experiments; see also Figs. 3D (main text) and SA4 and SA5. The inset in Fig. SB10 shows the same data for small densities and attachment rates. Note that the Figure may not display the steady-state values of the density, as the simulations converged only very slowly to the stationary state for high concentrations. The simulated time was *t* = 8 · 10^5^ s. The parameter values used in the simulation are listed in Table SB1. A lattice size of L = 2750 was used.

#### B9 Convolution method for image generation

To generate images from simulations that are comparable to images of the microscopy setup, we convoluted the particle occupation of the lattice with a point spread function: *I*(*r*) = *I*_0_ · (2 *J*_1_(2*πr*/*β*)/(2*πr*/*β*))^2^, with *r* – the radial distance to the origin, *I*_0_ – the maximal intensity, *J*_1_ – the Bessel function of the first kind of order one, and *β* = 0.78. The optical resolution of the experimental setup was estimated to be 400 nm, which is approximately reproduced by the chosen point-spread function. A plot of the corresponding function is shown in Fig. SB11.

**Figure SB11.**
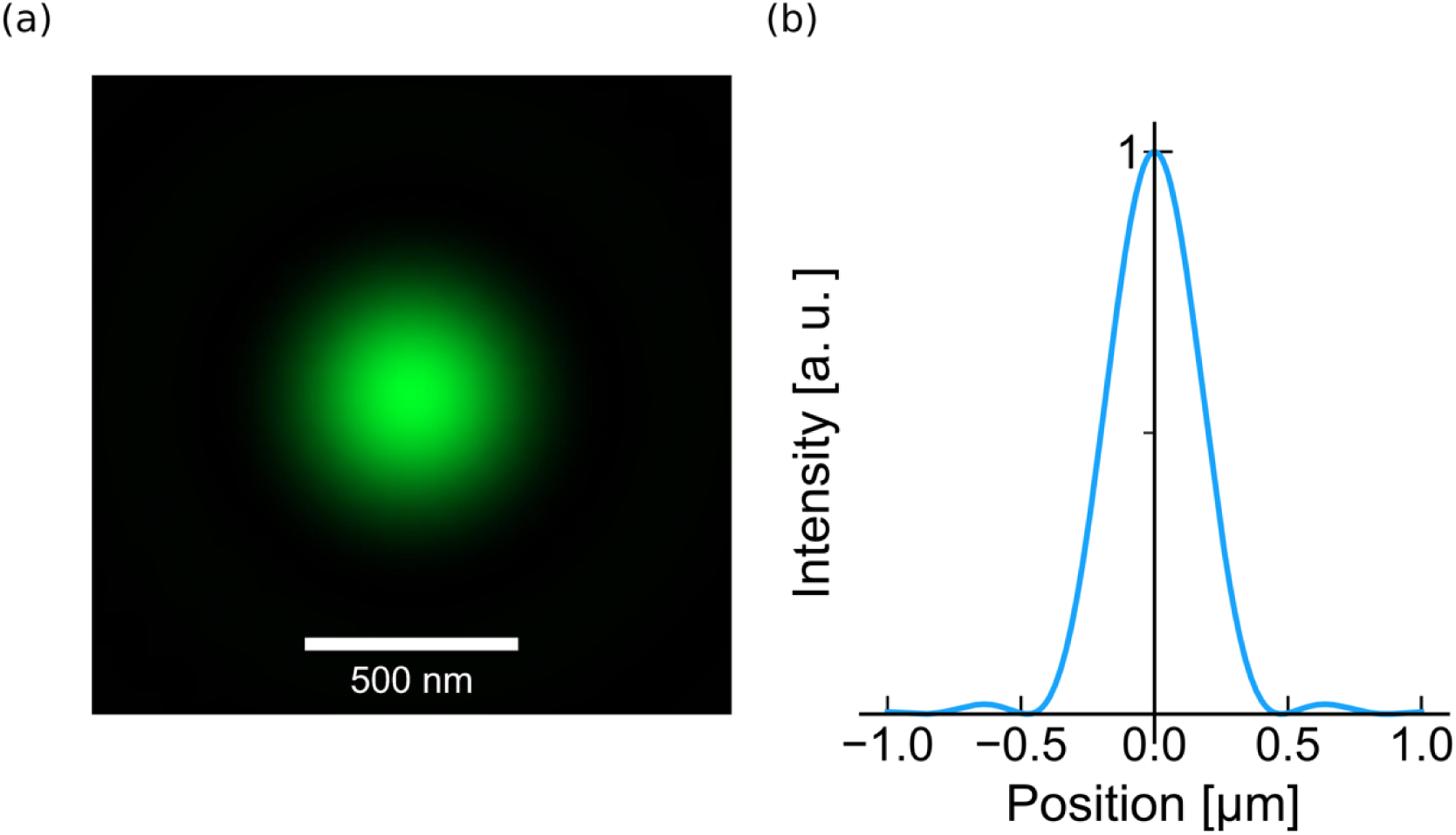
Illustration of the point-spread function used to generate images from the simulated kymographs. We used a point-spread function corresponding to an estimated optical resolution of approximately 400 nm. The function that was used to generate the kymographs is shown as a two-dimensional plot in **(a)** and the corresponding profile along the x-axis is shown in **(b)**.

## Footnotes

1 Here, the sign convention is such that positive signs denote attractive forces.

2 Note that—although originating from completely different processes—the proposed caterpillar mechanism for cluster motion is reminiscent of the inchworm mechanism that was speculated for the motion of the two heads of an individual kinesin: In both models, motion is the result of a wave-like motion of the different building blocks (a single kinesin head for the inchworm mechanism and an individual tetrameric motor for the caterpillar mechanism, respectively), with always the same building block leading

1 Here, we follow the convention to use positive forces for attractive interactions.

